# Powering electronic implants by high frequency volume conduction: in human validation

**DOI:** 10.1101/2021.03.15.435404

**Authors:** Jesus Minguillon, Marc Tudela-Pi, Laura Becerra-Fajardo, Enric Perera-Bel, Antonio J. del-Ama, Ángel Gil-Agudo, Álvaro Megía-García, Aracelys García-Moreno, Antoni Ivorra

**Affiliations:** Department of Information and Communications Technologies, Universitat Pompeu Fabra, 08018 Barcelona, Spain; Serra Húnter Fellow Programme, Universitat Pompeu Fabra, 08018 Barcelona, Spain; Department of Mathematics, Materials and Electronic Technology, Rey Juan Carlos University, 28933 Móstoles, Spain; Biomechanics and Technical Aids Department, National Hospital for Paraplegics (SESCAM), 45071 Toledo, Spain

**Keywords:** Volume conduction, Medical device, AIMD, Wireless power transfer, WPT, Galvanic coupling, First-in-human

## Abstract

Wireless power transfer (WPT) is frequently used as an alternative to batteries to accomplish miniaturization in electronic medical implants (eMIs). However, established WPT methods require bulky parts within the implant or cumbersome external systems, hindering minimally invasive deployments and the development of networks of eMIs. As an alternative, we propose a WPT approach based on volume conduction of high frequency (HF) current bursts. These currents are applied through external textile electrodes and are collected by the eMIs through two electrodes at their opposite ends. This approach avoids bulky components to obtain power, making it possible to develop implants with a flexible threadlike conformation. In here we study in humans if HF (6.78 MHz) current bursts complying with safety standards and applied through two textile electrodes strapped around a limb can provide substantial powers from pairs of implanted electrodes. Time averaged electric powers obtained from needle electrodes (diameter = 0.4 mm, length = 3 mm, separation = 30 mm) inserted into arms and lower legs of five healthy participants were 5.9 ± 0.7 mW and 2.4 ± 0.3 mW respectively. We also report a procedure to characterize the coupling between the external system and the implants using two-port impedance models generated from medical images. The results demonstrate for the first time in humans that innocuous and imperceptible HF current bursts that flow through the tissues by volume conduction can be used to wirelessly power threadlike eMIs, overcoming the limitations of existing WPT methods in terms of minimal invasiveness and usability.

## Introduction

Wireless power transfer (WPT) is frequently used in electronic medical implants as an alternative to electrochemical batteries. WPT offers two major advantages for the implant: longevity and miniaturization. In addition to electrochemical batteries and nuclear batteries^1^, other intrinsic power mechanisms have been explored such as those labeled as energy scavengers or energy harvesters^2–6^. However, all these alternative intrinsic power generation mechanisms also require voluminous parts (e.g. oscillating weights or thermopiles) to be integrated within the implants and provide powers considered to be insufficient for most implanted devices^7^.

To the best of our knowledge, the only WPT methods in clinical use are near-field inductive coupling and ultrasonic acoustic coupling^8^, the former being much more prevalent than the latter. Other WPT methods under exploration are: optic WPT^9–13^, mid-field inductive coupling^14,15^, far-field coupling^16^, capacitive coupling^17–19^ and WPT based on volume conduction which is also, less accurately, referred to as galvanic coupling^20–23^. Comprehensive and partial recent reviews on these methods can be found in^24–27^.

Compared to other WPT methods, galvanic coupling and capacitive coupling offer the advantage of not requiring a cumbersome external system or integrating bulky parts, such as piezoelectric crystals or coils, within the receiving implant for absorbing the energy transferred by the remote transmitter. The energy can be readily picked-up with a pair of thin electrodes separated by a few millimeters or centimeters. In addition, galvanic coupling and capacitive coupling are compatible with metallic hermetic packages (for housing the electronics) that can be made thinner than their glass or ceramic counterparts.

By using WPT based on high frequency (HF) volume conduction, we envision the development of injectable devices such as the one illustrated in Figure 1a. That is, we envision implantable devices with a diameter below 1 mm which consist of a very thin flexible body with two electrodes at opposite ends and a hermetic capsule with electronics within the flexible body. Because of their thinness and flexibility, it will be possible to implant these devices through minimally invasive procedures such as injection (as we have already demonstrated in^28^) or catheterization.

**Figure 1.**
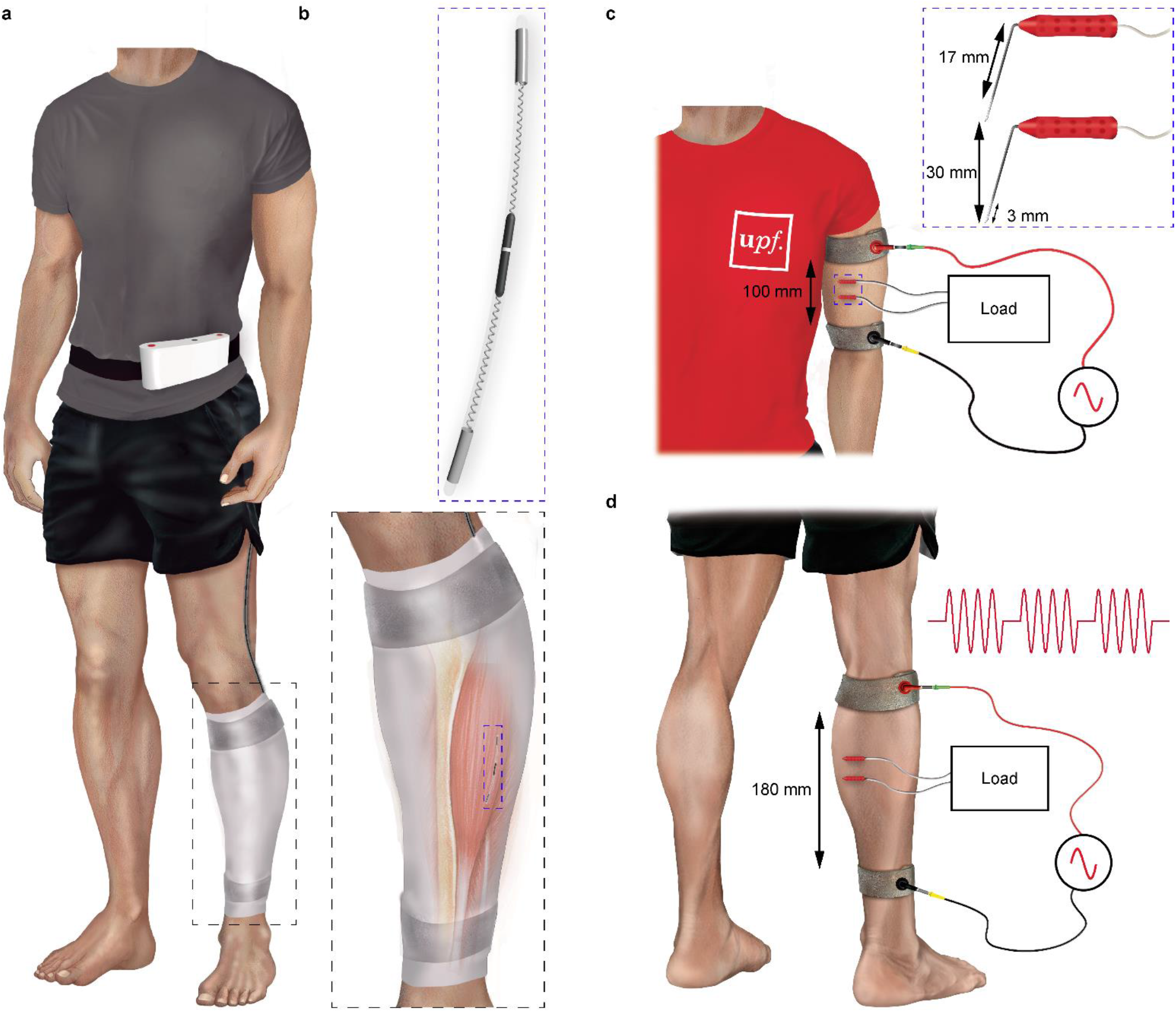
Hypothetical scenario of use for wireless power transfer (WPT) based on high frequency volume conduction and experimental setups employed in the present study. **a**, Hypothetical scenario: a threadlike implant (e.g. for neuromuscular electrical stimulation of the tibialis anterior in foot drop patients^45^) is powered by high frequency current bursts that are applied by a portable generator through a pair of external textile electrodes. **b**, Enlarged view of the implant, and its location in the tibialis anterior muscle. **c**, Upper limb (arm) experimental setup: an external electrical load (resistive) is connected to a pair of needle electrodes inserted into the brachial biceps. **d**, Lower limb (lower leg) experimental setup.

In the context of WPT for electronic medical implants, the distinction between galvanic coupling and capacitive coupling is fuzzy. In both cases the transmitter generates an alternating electric field across two electrodes and this field is picked-up by two electrodes of the receiver. The term galvanic coupling is generally employed when the transmitter electrodes and the receiver electrodes are directly in contact with the tissues (without any sort of insulation) and the frequencies are not very high (< 100 MHz). On the other hand, the term capacitive coupling is generally employed when the electrodes are coated with a dielectric and the frequencies are very high (> 100 MHz). Due to the passive electrical properties of living tissues, neither outright conductors nor outright dielectrics, in both cases the ac currents that flow through the tissues have a real component (corresponding to conduction currents, hence the term galvanic coupling) and an imaginary component (corresponding to displacement currents). The term volume conduction, therefore, is more accurate than the terms galvanic or capacitive coupling since both mechanisms are typically acting at the same time. We advocate for the use of frequencies in the order of some MHz or a very few tens of MHz to be able to power deep implants by avoiding the skin effect which in living tissues is very significant at frequencies above 100 MHz^29^. At the frequencies we propose, the conduction currents are substantially higher than the displacement currents and hence the use of the term galvanic coupling is justified^22^.

Remarkably, although galvanic coupling for intrabody communications has been studied lately by different research groups^30–32^ and it is even employed in implants in clinical use^33^, it appears that recently only in a very few occasions, besides in our own publications^20,22^, its use for powering implants has been explicitly proposed^21,23,34,35^. To this group can be added a few studies in which volume conduction at frequencies substantially below 100 MHz was proposed albeit referred to as capacitive coupling^18,36–39^. The absence of clinical systems using volume conduction for WPT is noteworthy, particularly taking into account that volume conduction for powering implanted devices was proposed more than 50 years ago^40^.We conjecture that such neglect mainly arises from not recognizing two crucial opportunities granted by volume conduction. First, large magnitude HF currents can safely flow through the human body if applied as short bursts (i.e. short enough to keep the overall power under the safety limit). Second, to obtain a sufficient voltage drop across its two intake (pick-up) electrodes, the implant can be shaped as a thin and flexible threadlike body (Figure 1b) which, as already stated, is a conformation suitable for minimally invasive deployment through injection or catheterization. Indeed, a singular feature of the conformation we propose to perform WPT based on volume conduction is the avoidance of flat structures for the receiver electrodes in the implant. Other researchers have typically proposed flat electrodes with a large surface area for maximizing power transfer efficiency, particularly in setups where the receiver electrodes are very close to the transmitter electrodes^35,41^. However, the use of large flat electrodes, albeit thin, precludes implantation via minimally invasive procedures and, therefore, results in systems which in terms of clinical applicability are similar to those based on inductive coupling. It can be stated that we trade power transfer efficiency off for minimal invasiveness.

The main goal of the present study is to illustrate that large magnitude HF currents can be innocuously and imperceptibly applied to humans and to demonstrate that those currents can produce substantial electric powers when picked-up by a pair of thin intramuscular electrodes separated a very few centimeters. The innocuity of the applied currents is ensured by adhering to available international safety standards for human exposure to electromagnetic fields^42,43^. In the present study, the IEEE standard^43^ was followed. The frequency of the applied sinusoidal currents (modulated as bursts) was set to 6.78 MHz because it corresponds to the central frequency of a designated ISM band^44^, thus minimizing the possibility of interfering radiocommunication systems, and because it is high enough to easily avoid risks related to unsought electrostimulation.

In the present study, in addition to reporting the results of the first in human validation of WPT based on volume conduction, we demonstrate a procedure to model this WPT method which characterizes coupling between the transmitters and the receivers by means of two-port impedance models which are generated from participants’ medical images (MRI). This procedure is not only relevant for the design of systems using WPT based on volume conduction but it could also be applied for modeling transmission channels in intrabody communications based on volume conduction.

## Results and discussion

### Maximum transferred power

Figure 2a shows a MRI image corresponding to the arm of participant P3. The markers of the external electrodes can be observed in this image. After building a 3D computational model from the MRI images, the electric field, the electric potential and the projected SAR (see definition in Subsection “Electrical safety” in “Methods”) distributions were numerically calculated. In this case (see Figure 2b) and in the other 9 cases (4 arms or A, 5 lower legs or LL; reported in Supplementary figure 14), it can be observed that, within the region encompassed by the two external electrodes, a few millimeters away from them, the electric field is coarsely uniform where the section of the limb smoothly changes. Consequently, the electric potential coarsely drops linearly between those electrodes (see Figure 2c and Supplementary figure 15). This is noteworthy for two reasons: 1) the electric field at the location of an implant (and hence the power obtained by the implant^22^) would not depend on the implantation depth, and 2) WPT based on volume conduction can be considered to be a non-focused WPT method since the applied electric field is present in the whole inter-electrode region. Non-focality implies that WPT based on volume conduction can be used for powering multiple implants with the same external energy source (this fact is illustrated in Subsection “Powering electronic devices” in “Results and discussion”). Examples of non-focused methods are those based on Helmholtz-like coil configurations^46^. Regarding the projected SAR distribution, it can be noted that maxima are generally located near the external electrodes. However, it is also worth noting that the values of these SAR maxima do not differ substantially from the SAR values where the needle electrodes were placed (see Figure 2d and Supplementary figure 13). Thus, the fact of having the SAR maxima close to the external electrodes does not significantly limit the maximum power transferred to the implants.

**Figure 2.**
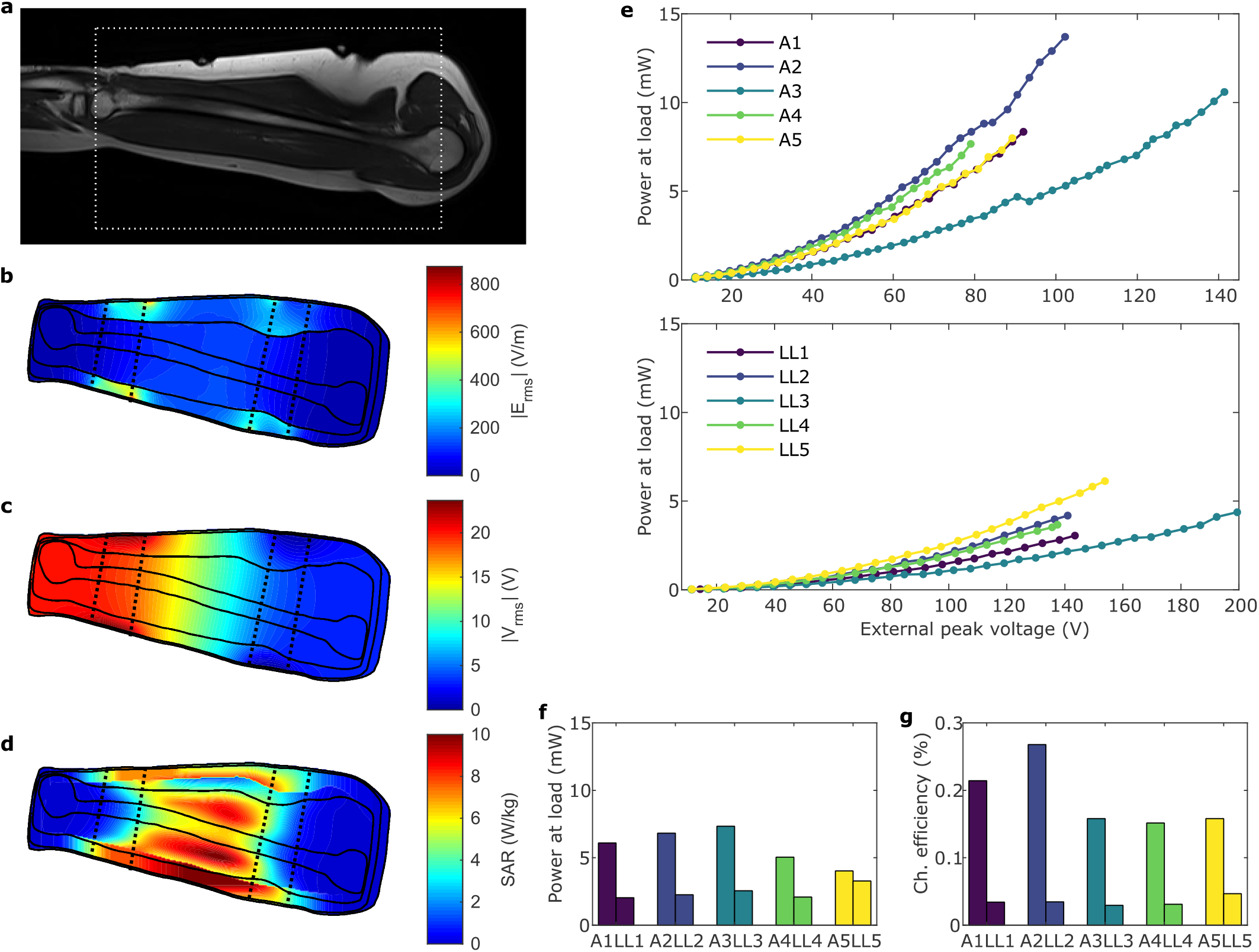
Examples of MRI, computed electric field, electric potential and projected SAR distributions, and power transfer experimental results. **a**, MRI image of the arm of participant P3 (i.e. A3). **b**, Computed electric field distribution inside A3. **c**, Computed electric potential distribution inside A3. **d**, Computed projected SAR distribution inside A3. **e**, Maximum transferred power to the optimal load connected across the pair of intramuscular needle electrodes when sinusoidal voltage bursts (carrier frequency = 6.78 MHz, burst duration = 100 μs and repetition frequency = 1 kHz) were applied across the pair of external electrodes: in the arms (i.e. A1 to A5) on top and in the lower legs (i.e. LL1 to LL5) on the bottom. Dots indicate experimental measurements. **f**, Maximum transferred power to the optimal load with a projected maximum SAR of 10 W/kg. **g**, Channel efficiency.

The average optimal load (i.e. the resistance that experimentally maximized the received ac power) was (244 ± 9) Ω (mean ± standard error of the mean, SEM) for the arms and (258 ± 6) Ω (mean ± SEM) for the lower legs (see Supplementary table 5 for details of the optimal load). The received ac powers (for the optimal load) for both limbs of all the participants are reported in Figure 2e. This time-averaged power was obtained in the load with it connected to the pair of intramuscular electrodes when sinusoidal voltage bursts (carrier frequency = 6.78 MHz, burst duration = 100 μs and repetition frequency = 1 kHz) were applied across the external electrodes. It can be observed that the received ac power shows a quadratic dependency on the externally applied voltage amplitude. Powers up to a projected maximum SAR of 20 W/kg, which is the limit imposed by the IEEE safety standard^43^ for limbs in controlled settings, are reported for arms of P2 and P5. In the remaining cases, the maximum reported power corresponds to a lower projected SAR due to technical limitations (i.e. maximum output amplitude of the generator) during the experimentation. For the arms, the maximum received power ranges from 7.7 mW (A4) to 13.7 mW (A2), with an average value of (10 ± 2) mW (mean ± SEM). For the lower legs, these values are noticeably lower. They range from 3.1 mW (LL1) to 6.1 mW (LL5), with an average value of (4.3 ± 0.6) mW (mean ± SEM). These results indicate that powers in the order of milliwatts can be transferred while complying with the safety standard.

For a projected maximum SAR of 10 W/kg, the received powers in the arms range from 4.0 mW (A5) to 7.3 mW (A3), with an average value of (5.9 ± 0.7) mW (mean ± SEM), and in the lower legs they range from 2.0 mW (LL1) to 3.3 mW (LL5), with an average value of (2.4 ± 0.3) mW (mean ± SEM) (see Figure 2f).

The channel efficiency for every single case is reported in Figure 2g. For the arms, it ranges from 0.15% (A4) to 0.27% (A2), with an average value of (0.19 ± 0.03)% (mean ± SEM). For the lower legs, it ranges from 0.029% (LL3) to 0.047% (LL5), with an average value of (0.035 ± 0.003)% (mean ± SEM). The difference between arms and lower legs is mainly due to the anatomical characteristics of both limbs (e.g. volume, section, length, fat thickness, etc.)^47^. These power transfer efficiencies in the order of 0.1% are much lower than those typically reported, in the order of 1% or even in the order of 10%, for WPT systems based on inductive coupling or ultrasonic acoustic coupling^26,48^. However, comparison in terms of efficiency with published WPT systems is not straightforward as efficiency depends on multiple factors such as the geometry of the elements (e.g. the dimensions of the external applicator and of the receiver) and the relative conformation of the elements of the system (e.g. whether the receivers are within a region encompassed by the applicator of the transmitter). In fact, for some conformations, volume conduction is advocated by some researchers because of its superior efficiency^26^. Comparison of the results reported here for the arm (efficiency around 0.2%) with those reported for a similar scenario in which inductive coupling was assayed *ex vivo* with an efficiency of around 2%^19^ suggests that volume conduction is ten times less efficient than inductive coupling. Nevertheless, this comparison must be considered as inconclusive because the receiver inductor of that study had a diameter of 4 mm and inductive coupling efficiency strongly depends on such diameter^49^. Further research is required regarding this aspect. In any case, the efficiency of volume conduction does not jeopardize the feasibility of using a wearable external system (such as the one showed in the envisioned scenario in Figure 1a). Taking into account the mean of the channel efficiency (i.e. 0.19% and 0.035% for arms and lower legs, respectively) and the power consumption of electronic medical implants (typically 1 mW or less), the externally applied power should be in the order of 0.5 and 3 W for the arms and lower legs, respectively. In the market, there are several wearable rechargeable batteries (e.g. 100 cm^3^ or less) that provide more than 30 W for one hour (e.g. 2447-3034-20-520 from Ansmann AG, Assamstadt, Germany).

### Experimental measurements versus computational model

The transmission channel formed by the band electrodes, the tissues and the needle electrodes was modeled as a two-port impedance network (see Figure 3a)^50^. The impedance (Z) parameters of the network were numerically computed using the 3D computational model of the limb obtained after segmenting the MRI images. The obtained two-port impedance networks allow simulating power transfer with reasonable accuracy. As an example, when applying a peak voltage of 79 V across the external electrodes, the average relative error between the experimentally received power and the simulated one is 4%, with a standard deviation of 26% (see Figure 3b-f). After analyzing the impact of different possible sources of error in the experimental procedures and in the simulations, it was determined that the variability in the insertion angle of the needle electrodes was the most likely cause of the discrepancies observed between the simulated and the experimental results. This variability causes significant random errors in the separation distance between the conductive tips of the electrodes. Even though the location of the insertion point of the needles was carefully ensured by using markers, the insertion was performed manually orthogonally to the skin without any provision to ensure the penetration angle. Simulations were performed assuming ±10° errors in the penetration angle. Considering this penetration angle error margin, the simulations fit the experimental results in all cases except for the arm of P2. This inconsistency may be caused by the fact that, in this case, the insertion marks of the needle electrodes had to be displaced 5 mm from the original position to avoid a blood vessel that was detected in the MRI images during the experimentation. Although the position of the needles was accordingly modified on the numerical model, this adjustment may explain the inconsistency.

**Figure 3.**
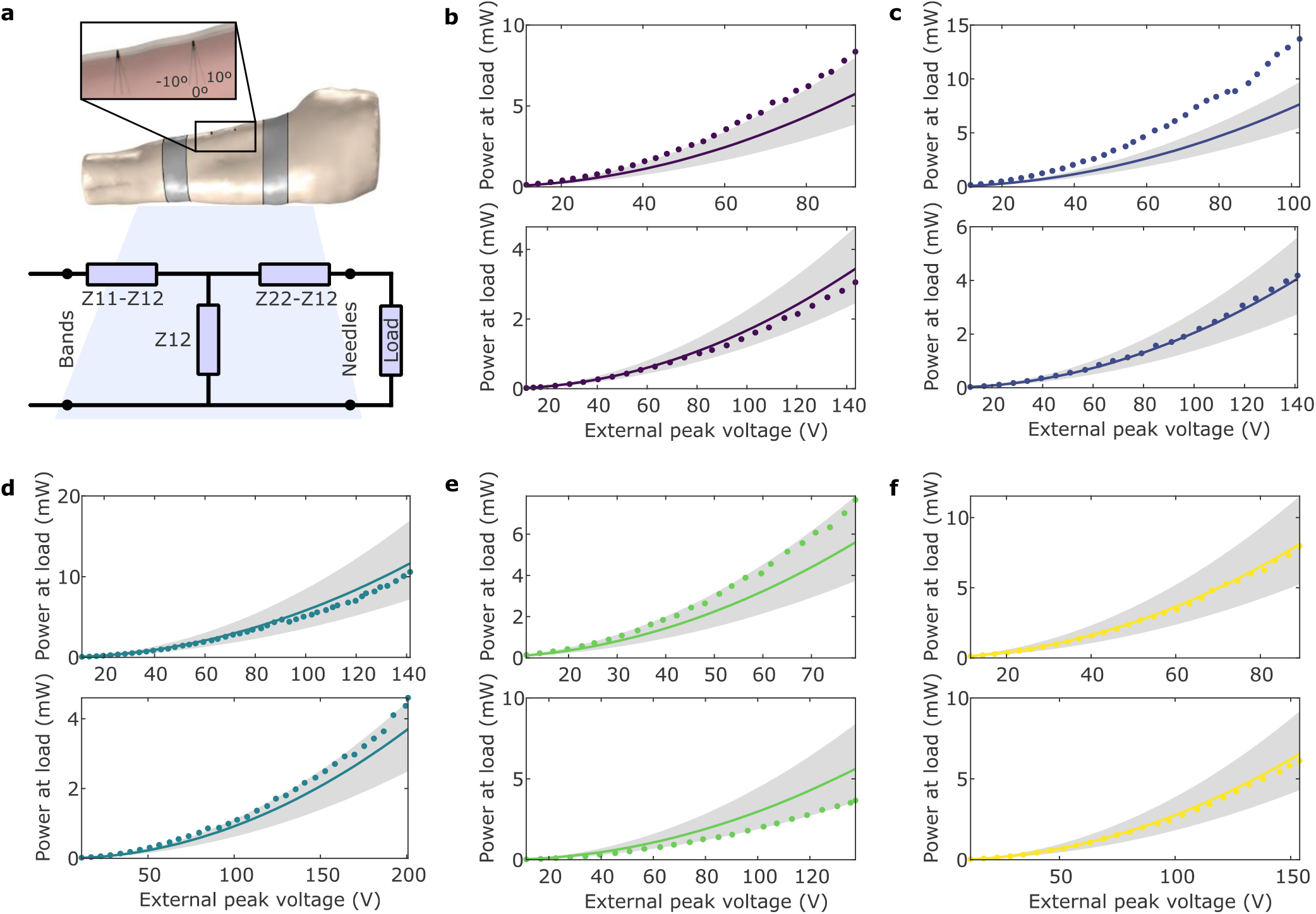
Computational modeling results. **a**, The transmission channel (i.e. limb tissues, band electrodes and needle electrodes) was modeled as a two-port network. The inlet shows the considered insertion angle error of the needle electrodes. **b**, Comparison of experimental maximum transferred power and simulated transferred power using the optimal experimental load in the arm (top) and in the lower leg (bottom) of P1. Dots indicate experimental measures. Solid line indicates simulated results under the assumption that that both needle electrodes are perfectly aligned (i.e. 0°). Upper shadow edge indicates simulated results under the assumption that both needle electrodes have an inclination of −10° (i.e. 10° towards the center). Lower shadow edge indicates simulated results under the assumption that both needle electrodes have an inclination of +10° (i.e. 10° towards the band electrodes). **c**, Same as Figure 3b for P2. **d**, Same as Figure 3b for P3. **e**, Same as Figure 3b for P4. **f**, Same as Figure 3b for P5.

### Powering electronic devices

Figure 4a shows a picture of an electronic device connected to the needle electrodes of the arm of P1, being safely powered according to the IEEE standard through volume conduction of HF current bursts (carrier frequency = 6.78 MHz, burst duration = 100 μs, and repetition frequency = 1 kHz). The electronic device connected to the needle electrodes does not contain any power source and it is composed by: 1) a circuit consisting of a bridge rectifier with dc blocking capacitors, a voltage regulator and a microcontroller and 2) an LCD screen (see insert in Figure 4a). These electronics are similar to those of most implantable medical devices in terms of complexity and power consumption (approximately 1 mW). Figure 4c reports the projected maximum SAR that appears in tissues when the amplitude of the external voltage is sufficient for powering the device in all participants and limbs. In all cases, the SAR is below 10 W/kg. It ranges from 2.3 W/kg (A3) to 4.0 W/kg (A5) for the arms, with an average value of (3.2 ± 0.4) W/kg (mean ± SEM). For the lower legs, it ranges from 6.2 W/kg (LL5) to 8.1 W/kg (LL1), with an average value of (7.0 ± 0.4) W/kg (mean ± SEM). Figure 4b shows two electronic devices being powered by the same pair of external electrodes in the lower leg of P4. Each device only draws a small portion of the total external energy without significantly distorting the electric field at the location of the other implant, thus enabling the possibility of increasing the total efficiency of the method by increasing the number of devices in the same powered area. This non-focused character of volume conduction makes it advantageous with respect to other focused WPT methods such as ultrasonic acoustic coupling.

**Figure 4.**
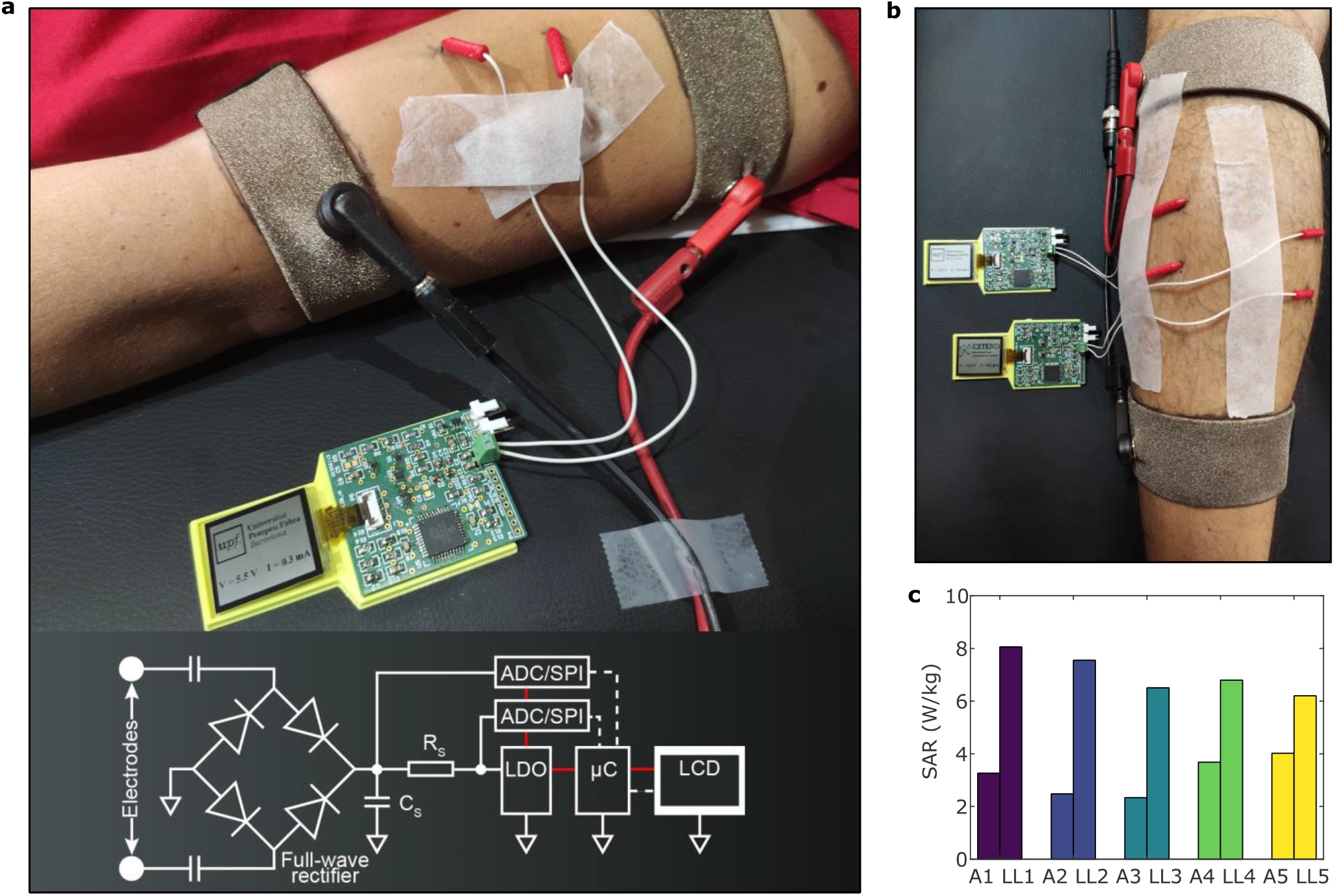
Illustration of the capability of WPT based on volume conduction for safely powering electronic devices. **a**, Picture of an electronic device, which is comparable to a medical implant in terms of circuit complexity and power consumption, being powered by volume conduction of high frequency current bursts (carrier frequency = 6.78 MHz, burst duration = 100 μs, and repetition frequency = 1 kHz). The insert shows the circuit diagram of the electronic device. **b**, Two devices being simultaneously powered by the same current bursts. **c**, Projected maximum SAR in tissues when the amplitude of the external voltage is sufficient for powering the electronic devices of this setup.

### 18-minute temperature experiment

This experiment was conducted to illustrate thermal safety ensured by the IEEE standard. That is, if the SAR is kept below limits, temperature increase in tissue will be very moderate -and thus harmless- even for prolonged exposures. The same HF current bursts (carrier frequency, burst duration, burst repetition frequency and voltage amplitude) that were used for powering the electronic device of the previous subsection were continuously applied on the lower leg of P4. After 18 minutes, the temperature did not noticeably increase near the electrodes (0.2 °C and 0.3 °C) or on the skin surface between them. In the contralateral limb, used as control with unpowered external electrodes (i.e. no burst applied), the temperature near the electrodes decreased slightly (0.3 °C and 0.4 °C). These small temperature variations, which in addition were not monotonic, may be mainly related to the fact of having the textile bands attached to the surface of the limb.

### Perception of HF current bursts and skin appearance

None of the participants reported any sensation related to heat or electrostimulation during the experimentation in both limbs. They barely reported uncomfortable sensations related to the fact of having the needles inserted. Therefore, the applied HF current bursts are not only innocuous, as guaranteed by adherence to the IEEE standard, but also imperceptible. No skin alterations were observable.

This study focuses on the safety of the method by following the IEEE standard on safety levels with respect to human exposure to electromagnetic fields. These aspects can be extrapolated to chronic implants but, for the final application (chronic use with implantable devices), there are other safety aspects (e.g. long-term electrode changes, mechanical stability, etc.) that should be analyzed in chronic animal studies.

## Conclusion

This study in healthy participants demonstrates that, albeit with poor efficiency, ac and dc electric powers in the order of milliwatts can be obtained from pairs of thin electrodes within limb muscles when HF sinusoidal current bursts are safely delivered through two textile electrodes shaped as bands strapped around the limb and encompassing the region where the pair of thin electrodes is located. In addition, it is demonstrated that these currents are imperceptible and that the obtained power from the pair of thin electrodes can be used to power complex electronic circuits with digital and analog functionalities. To the best of our knowledge, this is the first time that the use of WPT based on HF volume conduction has been validated in humans. Since none of the observations and principles preclude the use of this approach in other comparable conditions (e.g. different waveforms, tissues, anatomical locations or geometries for the systems), the results of this study pave the road for the development of diagnostic and therapeutic systems using threadlike electronic implants powered by WPT based on HF volume conduction in limbs.

In addition to unprecedented minimal invasiveness, other remarkable advantages of the presented approach over other WPT methods are the capabilities to: 1) power deep implants, 2) simultaneously power multiple implants with the same external applicator, 3) deliver high peak powers and 4) avoid inconvenient external applicators and elements such as rigid bulky coils for inductive coupling or gels for ultrasonic acoustic coupling.

Furthermore, this study proposes and demonstrates a procedure to accurately model WPT based on volume conduction that characterizes the coupling between the transmitters and the receivers by means of two-port impedance models generated from medical imaging data. This procedure is not only relevant for the design of systems using WPT based on volume conduction but it could also be applied for modeling transmission channels in intrabody communications based on volume conduction.

## Methods

### Participants

Five young (age range from 21 to 38 years) healthy volunteers participated in the study: P1 (female), P2 (female), P3 (female), P4 (male) and P5 (male). They were recruited through a call for participation sent by email to colleagues and were not paid for their participation. Before starting the experimental procedure, they were provided with oral and written information regarding the study (including risks, benefits and data protection aspects) and signed an informed consent form. The study was conducted in the National Hospital for Paraplegics in Toledo (Spain). The experimental protocol was approved by the Ethics Committee for Clinical Investigation of the Complejo Hospitalario de Toledo (December 5, 2019; reference number: 467).

### Experimental procedures

The experimental procedures were divided into two phases: 1) magnetic resonance imaging (MRI) images acquisition and 2) assays. During the first phase, MRI images were acquired of the non-dominant upper (arm) and lower (leg) limb of each participant. These images were used to build 3D computational models for numerical calculations (see Subsection “MRI acquisition and segmentation” in this section). Prior to MRI acquisition, the planned locations for the external and for the intramuscular electrodes were marked with MRI fiducial markers (see Supplementary method 1 for details of the marking procedure). The position of the participants during MRI acquisition was the same as that during the second phase of the experimental procedures. Once all the participants finished the first phase, they sequentially participated in the second phase. During the second phase, different procedures and assays were conducted. First, the participant was positioned on a stretcher (supine and prone positions for arm and lower leg, respectively) and instructed to avoid unnecessary movements. A pair of external electrodes was strapped around the corresponding limb. As control, another pair of electrodes was strapped around the contralateral limb for later discerning whether skin alterations could be due to the delivery of the HF currents or were caused by the materials of the electrodes. These external electrodes consisted in textile bands with a width of 3 cm (used for arms) or 4 cm (used for lower legs) and were made of conductive fabric. The electrodes on the non-dominant limb were connected to a HF voltage generator (see Supplementary method 2 for details of the external electrodes and the generator). Sinusoidal voltage bursts (carrier frequency of 6.78 MHz and FB = 0.1; where F = 1 kHz is the repetition frequency of the bursts and B = 100 μs is their duration) of different amplitudes (increasing order) were applied to check if the participant perceived some discomfort. Once the preliminary assay was performed (without any notification of sensations by any of the participants), a pair of intramuscular needle electrodes were inserted either into the brachial biceps or into the medial gastrocnemius of the non-dominant limb for the arm and lower leg, respectively, under aseptic conditions. These needle electrodes had a diameter of 0.4 mm and a length of 20 mm of which only the distal 17.5 mm were inserted. The needle has a 3 mm long exposed surface on its tip (530607 from Inomed Medizintechnik GmbH, Emmendingen, Germany; see Figure 1). The needle electrodes were connected to a discrete potentiometer (3683S-1-202L from Bourns Inc., Riverside, CA, US) using short cables (maximum of 12 cm) to avoid inductive and capacitive wiring effects. After that, the remaining assays (i.e. maximum transferable power, and screen power-up, and, in one case, temperature evolution in 18 minutes) were conducted. Once the last assay was finished, the textile electrodes were unstrapped, the needle electrodes were extracted and the skin areas where the electrodes had been located were inspected. No damage was observed in these areas or in the electrodes (see Supplementary figures 16 and 17). Only one case (arm of P2) presented small-sized superficial hematomas in the areas where the needle electrodes had been inserted approximately one hour after the extraction. The temperature of the limb was monitored using a thermal imaging infrared camera (see Supplementary method 3 for details of the infrared camera) during the entire phase. The second phase was repeated for the other limb. The order of the limbs was randomized among the participants.

### MRI acquisition and segmentation

Each participant underwent MRI images acquisition, recording four sequences per limb: T1 axial (slice resolution = 0.52 × 0.52 mm; slice thickness = 6 mm), T1 sagittal (slice resolution = 1.04 × 1.04 mm; slice thickness = 4.8 mm), T1 coronal (slice resolution = 1.04 × 1.04 mm; slice thickness = 4.65 mm), and T2 axial (slice resolution = 0.52 × 0.52 mm; slice thickness = 6 mm). These sequences were acquired with a 3 tesla system (Magnetom Trio, a Tim System from Siemens Healthcare GmbH, Erlangen, Germany). Four different tissues were segmented: 1) bone, which included cortical and trabecular tissues, bone marrow, and articular cartilages (e.g. meniscus), 2) muscle, including tendons, ligaments, and intramuscular fat, 3) subcutaneous fat, and 4) skin. For the segmentation of bone, muscle and fat, the procedure started by manually segmenting the T1 axial acquisition every other slice. The initial segmentation of the skin had to be performed using a different procedure since it was not visible in the entire MRI stack because of its thinness. The skin thickness was approximated by measuring it in different slices of the MRI, and then the skin was generated by performing a 2D axial dilation on the fat segmentation. The measured skin thicknesses are reported in Supplementary table 6. For each tissue, the remaining slices were interpolated with the “3D interpolation” built-in tool of The Medical Imaging Interaction Toolkit from the German Cancer Research Center, Heidelberg, Germany. This interpolation was based on the radial basis function interpolation^51^ and on Laplacian smoothing^52^. Minor manual adjustments were required to correct sharp curvatures of the geometry in the interpolated slices. Finally, smoothed surface meshes were generated from the 3D interpolation (see Supplementary figure 18) and then exported for numerical computation. The MRI fiducial markers (three for each band electrode and one for each needle electrode) were located in all three orthogonal stacks (i.e. axial, sagittal and coronal) and their center was precisely annotated.

### Assay for measuring the maximum transferable power

The maximum received power for an optimal resistive load was obtained. For that, first, sinusoidal voltage bursts (carrier frequency of 6.78 MHz and FB = 0.1 (F = 1 kHz)) were applied across the external electrodes and the optimal load (i.e. the resistance that provides maximum power transfer) was experimentally found by adjusting the resistance of the discrete potentiometer. Then, with the optimal load connected to the intramuscular electrodes and the same voltage waveform applied across the external electrodes, different amplitudes were applied. The externally applied voltage amplitude and current amplitude, together with the voltage amplitude at the load (*V*_*load*_), were measured (see Supplementary method 3 for details of the measurement apparatus). For each amplitude, the received power at the load was computed as 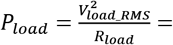 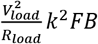, where *R*_*load*_ is the optimal load and the scaling factor 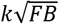, with 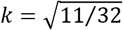, transforms amplitude values into RMS values for the applied waveform (see Supplementary method 2 for details of the applied waveform). The average channel efficiency was also computed as the ratio between the received power and the externally applied power.

### Screen power-up assay

The potentiometer was replaced by an electronic device consisting of a circuit that did not contain any power source and that was connected to an LCD screen (see insert in Figure 4 and Supplementary method 4 for details of the demonstrative electronic device). In terms of complexity and power consumption (approximately 1 mW), this device is comparable to the electronics of most implantable electronic devices, and far above in terms of power consumption to that of pacemakers, which is in the order of 10 μW ^3^. The assay consisted in finding out the required external voltage amplitude to power-up the electronic device. With the same external HF voltage waveform as above (carrier frequency of 6.78 MHz and FB = 0.1 (F = 1 kHz)), the external amplitude was increased until the screen powered-up and displayed information. Both the externally applied voltage and current were measured. In addition, the dc voltage and the dc current were measured at the load. In one case (lower leg of P4), for illustrating the capability of the proposed approach to power multiple devices, a second identical device was connected to an additional pair of electrodes inserted in the lateral gastrocnemius of the same limb (Figure 4).

### Temperature evolution in 18 minutes

In one case (lower limb of P4), the same external HF voltage waveform as above (carrier frequency of 6.78 MHz and FB = 0.1 (F = 1 kHz)) with fixed amplitude (the same as for powering-up the screen) was uninterruptedly applied for 18 minutes. This was performed once the intramuscular electrodes had been extracted. With the participant seated in a chair, the temperature variations were monitored and recorded using the infrared camera. The contralateral limb was used as control: external electrodes were strapped around the lower leg, but no voltage was applied.

### Electrical safety

Safety standards for human exposure to electromagnetic fields^42,43^ identify two general sources of risk regarding passage of alternating currents through the body which limit the amplitude of the currents that can be applied: 1) risks caused by unsought electrical stimulation of excitatory tissues and 2) risk of thermal damage due to Joule heating. Risks caused by unsought stimulation can be avoided if the frequency of the applied currents is high enough. In particular, the standard followed in the present study (the IEEE standard^43^) determines that, for continuous sinusoidal currents with a frequency above 5 MHz, electrostimulation is not a matter of concern. The electrostimulation risks are especially relevant for low frequencies (< 100 kHz). However, if the sinusoidal currents are applied in the form of bursts, the generated low frequency harmonics must be considered^22^. In this study, sinusoidal voltage bursts (carrier frequency of 6.78 MHz and FB = 0.1 (F = 1 kHz)) were applied. To minimize the contribution of the generated low frequency harmonics, the applied bursts were smoothed with a tapered cosine window (with *r = 0.5*, see Supplementary method 2 for details of the applied waveform). Applying the expression for non-sinusoidal fields established in the safety standard to the used voltage waveform, the maximum peak electric field that can be applied (i.e. the maximum *in situ* electric field to avoid electrostimulation) is above 200 MV/m (see Supplementary method 5 for details of the calculation). This limit is far above the electric field amplitudes that were computed to be produced during the experimental sessions. Risk of thermal damage due to Joule heating is addressed by the standards by imposing a limitation to the so-called specific absorption rate (SAR) which has units of W/kg and indicates the heat dissipated per unit of tissue mass due to Joule heating. In the case of limbs, the IEEE standard indicates that the maximum admissible SAR is 20 W/kg, space-averaged over any cubical 10 g of tissue and time-averaged for 6 minutes. In order to ensure that the SAR restriction was met during the experimentation, the external voltages were applied for short time exposures (i.e. < 30 seconds). On the other hand, the SAR values indicated in Section “Results and discussion” correspond to *projected SAR* values for exposures longer than 6 minutes. That is, the electric field magnitudes obtained with the 3D computational models (see Subsection “Numerical methods” in this section) were used to compute the SAR that would be produced by sinusoidal voltage bursts continuously applied for 6 minutes or more. The IEEE standard indicates another limit with respect to the whole body SAR averaged for 1 hour. In all cases, the projected whole body SAR was below this limit (i.e. 0.4 W/kg) for exposures longer than 1 hour.

### Perception of heat-related and electrical-stimulation-related sensations

During the second phase of the experimental procedures, self-perception of heat-related (HR) and electrical-stimulation-related (ESR) sensations was monitored. In particular, the participants were asked for oral notification of any HR or ESR sensation at any time without knowing when the HF voltage bursts were applied. In case of sensation, the HR perception was classified into three intensity levels (*1-Not sure*, *2-Pleasant warmth* and *3-Unpleasant warmth*) and the ESR perception was classified into three intensity levels (*1-Not sure*, *2-Pleasant sensation* and *3-Unpleasant sensation*) and five categories (*1-Tingling*, *2-Puncture*, *3-Pressure*, *4-Pain* and *5-Other*).

### Numerical methods

The meshes obtained from the MRI segmentation procedure were imported into COMSOL Multiphysics 5.3 (from COMSOL, Inc., Burlington, MA, US) to create a 3D computational model of each limb. This model included four tissue types (skin, subcutaneous fat, muscle and bone). The passive electrical properties and densities of the tissues are reported in Supplementary table 1. The needle electrodes (modeled as cylinders) and the external electrodes (modeled as superficial bands wrapping the tissues) were added to the 3D computational model. The electrical properties of the electrodes are reported in Supplementary table 2. The transmission channel formed by the band electrodes, the tissues and the needle electrodes was modeled as a two-port impedance network^50^. The impedance (Z) parameters of the network were numerically computed using the 3D computational model simulating the delivery of ac currents (frequency = 6.78 MHz) either through the external electrodes or through the needle electrodes. These values are reported in Supplementary tables 3 and 4. In addition, the 3D computational model was used to calculate the electric potential and the electric field magnitude distributions inside the tissues. For that, a sinusoidal voltage (frequency = 6.78 MHz) was applied to the external electrodes, being its RMS value equal to the RMS value of the experimental applied waveform. The transformation from amplitude values into RMS values was done by applying the mentioned scaling factor 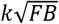, with 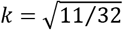. The projected local SAR was calculated from the electric field magnitude according to the safety standard, with equation 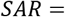 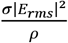, where *σ* is the local tissue conductivity, *ρ* is the local tissue density and *E*_*rms*_ is the RMS value of the computed electric field. The SAR was space-averaged over any cubical 10 g of tissue following the guidelines of the standard^43,53^. See Supplementary method 5 for details of the numerical methods.

## Supporting information

Supplementary material (MATLAB scripts)

## Acknowledgements

This work was supported by the European Union’s Horizon 2020 Research and Innovation Programme under Grant 779982. AI gratefully acknowledges the financial support by ICREA under the ICREA Academia Programme.

## Author contributions

A.I. conceived and supervised the study. J.M., M.T.P., L.B.F. and A.I. designed the experimental protocols. J.M., L.B.F., A.J.A., A.G.A., A.M.G. and A.I. prepared the documents for the Ethics Committee. J.M. and M.T.P implemented the devices. A.G.A. was the physician in charge of the medical procedures. J.M., M.T.P. L.B.F., E.P., A.J.A., A.M.G. and A.G.M. performed the study. E.P. performed the MRI segmentation. J.M. and M.T.P. conducted the analysis and interpreted the data. L.B.F. and A.I. reviewed analyses and results. J.M., M.T.P., L.B.F. and A.I. wrote the manuscript. All authors reviewed and edited the manuscript.

## Additional information

### Supplementary information

Supplementary information is available for this paper.

### Code availability

MATLAB scripts for computing the SAR and the maximum *in situ* electric field to avoid electrostimulation of the IEEE safety standard are available as Supplementary material.

## References

1. Parsonnet, V., Driller, J., Cook, D. & Rizvi, S. A. Thirty-one years of clinical experience with nuclear-powered pacemakers. PACE - Pacing Clin. Electrophysiol. 29, 195–200 (2006).

2. Han, M. et al. Three-dimensional piezoelectric polymer microsystems for vibrational energy harvesting, robotic interfaces and biomedical implants. Nat. Electron. 2, 26–35 (2019).

3. Cadei, A., Dionisi, A., Sardini, E. & Serpelloni, M. Kinetic and thermal energy harvesters for implantable medical devices and biomedical autonomous sensors. Meas. Sci. Technol. 25, 012003 (2014).

4. Nadeau, P. et al. Prolonged energy harvesting for ingestible devices. Nat. Biomed. Eng. 1, 1–8 (2017).

5. Jiang, D. et al. A 25-year bibliometric study of implantable energy harvesters and self-powered implantable medical electronics researches. Mater. Today Energy 16, 100386 (2020).

6. Hinchet, R. et al. Transcutaneous ultrasound energy harvesting using capacitive triboelectric technology. Science (80-. ). 365, 491–494 (2019).

7. Thimot, J. & Shepard, K. L. Bioelectronic devices: Wirelessly powered implants. Nat. Biomed. Eng. 1, 1–2 (2017).

8. Auricchio, A. et al. Feasibility, safety, and short-term outcome of leadless ultrasound-based endocardial left ventricular resynchronization in heart failure patients: Results of the Wireless Stimulation Endocardially for CRT (WiSE-CRT) study. Europace 16, 681–688 (2014).

9. Gutruf, P. et al. Fully implantable optoelectronic systems for battery-free, multimodal operation in neuroscience research. Nat. Electron. 1, 652–660 (2018).

10. Abdo, A. et al. Floating light-activated microelectrical stimulators tested in the rat spinal cord. J. Neural Eng. 8, (2011).

11. Kim, J. et al. Active photonic wireless power transfer into live tissues. Proc. Natl. Acad. Sci. U. S. A. 117, 16856–16863 (2020).

12. Mathieson, K. et al. Photovoltaic retinal prosthesis with high pixel density. Nat. Photonics 6, 391–397 (2012).

13. Zhang, H. et al. Wireless, battery-free optoelectronic systems as subdermal implants for local tissue oximetry. Sci. Adv. 5, eaaw0873 (2019).

14. Ho, J. S. et al. Wireless power transfer to deep-tissue microimplants. Proc. Natl. Acad. Sci. U. S. A. 111, 7974–7979 (2014).

15. Agrawal, D. R. et al. Conformal phased surfaces for wireless powering of bioelectronic microdevices. Nat. Biomed. Eng. 1, 1–9 (2017).

16. Chow, E. Y., Ouyang, Y., Beier, B., Chappell, W. J. & Irazoqui, P. P. Evaluation of cardiovascular stents as antennas for implantable wireless applications. IEEE Trans. Microw. Theory Tech. 57, 2523–2532 (2009).

17. Jegadeesan, R., Agarwal, K., Guo, Y. X., Yen, S. C. & Thakor, N. V. Wireless Power Delivery to Flexible Subcutaneous Implants Using Capacitive Coupling. IEEE Trans. Microw. Theory Tech. 65, 280–292 (2017).

18. Erfani, R., Marefat, F., Sodagar, A. M. & Mohseni, P. Transcutaneous capacitive wireless power transfer (C-WPT) for biomedical implants. Proc. - IEEE Int. Symp. Circuits Syst. 8–11 (2017). doi:10.1109/ISCAS.2017.8050940

19. Aldaoud, A. et al. Near-field wireless power transfer to stent-based biomedical implants. IEEE J. Electromagn. RF Microwaves Med. Biol. 2, 193–200 (2018).

20. Becerra-Fajardo, L., Schmidbauer, M. & Ivorra, A. Demonstration of 2 mm Thick Microcontrolled Injectable Stimulators Based on Rectification of High Frequency Current Bursts. IEEE Trans. Neural Syst. Rehabil. Eng. 25, 1343–1352 (2017).

21. Chen, P., Yang, H., Luo, R. & Zhao, B. A Tissue-Channel Transcutaneous Power Transfer Technique for Implantable Devices. IEEE Trans. Power Electron. 33, 9753–9761 (2018).

22. Tudela-Pi, M., Becerra-Fajardo, L., Garcia-Moreno, A., Minguillon, J. & Ivorra, A. Power Transfer by Volume Conduction: In Vitro Validated Analytical Models Predict DC Powers Above 1 mW in Injectable Implants. IEEE Access 8, 37808–37820 (2020).

23. Sedehi, R. et al. A Wireless Power Method for Deeply Implanted Biomedical Devices via Capacitively Coupled Conductive Power Transfer. IEEE Trans. Power Electron. 36, 1870–1882 (2021).

24. Amar, A. Ben, Kouki, A. B. & Cao, H. Power approaches for implantable medical devices. Sensors (Switzerland) 15, 28889–28914 (2015).

25. Kim, A., Ochoa, M., Rahimi, R. & Ziaie, B. New and Emerging Energy Sources for Implantable Wireless Microdevices. IEEE Access 3, 89–98 (2015).

26. Barbruni, G. L., Motto Ros, P., Demarchi, D., Carrara, S. & Ghezzi, D. Miniaturised Wireless Power Transfer Systems for Neurostimulation: A Review. IEEE Trans. Biomed. Circuits Syst. 1–20 (2020). doi:10.1109/tbcas.2020.3038599

27. Agarwal, K., Jegadeesan, R., Guo, Y. X. & Thakor, N. V. Wireless Power Transfer Strategies for Implantable Bioelectronics. IEEE Rev. Biomed. Eng. 10, 136–161 (2017).

28. Ivorra, A., Becerra-Fajardo, L. & Castellví, Q. In vivo demonstration of injectable microstimulators based on charge-balanced rectification of epidermically applied currents. J. Neural Eng. 12, 066010 (2015).

29. Vander Vorst, A., Rosen, A. & Kotsuka, Y. RF/Microwave Interaction Mechanisms in Biological Materials. in RF/Microwave Interaction with Biological Tissues (John Wiley & Sons, Inc., 2006).

30. Tomlinson, W. J., Banou, S., Blechinger-Slocum, S., Yu, C. & Chowdhury, K. R. Body-guided galvanic coupling communication for secure biometric data. IEEE Trans. Wirel. Commun. 18, 4143–4156 (2019).

31. Seyedi, M., Kibret, B., Lai, D. T. H. & Faulkner, M. A survey on intrabody communications for body area network applications. IEEE Trans. Biomed. Eng. 60, 2067–2079 (2013).

32. Lindsey, D. P., McKee, E. L., Hull, M. L. & Howell, S. M. A new technique for transmission of signals from implantable transducers. IEEE Trans. Biomed. Eng. 45, 605–613 (1998).

33. Sharma, D. et al. The leadless cardiac pacemaker conductive communication. JACC Clin. Electrophysiol. 1, 335–336 (2015).

34. Zhu, W. et al. Volume conduction energy transfer for implantable devices. J. Biomed. Res. 27, 509–514 (2013).

35. Hackworth, S. A. Design, Optimization, and Implementation of a Volume Conduction Energy Transfer Platform For Implantable Devices. University of Pittsburgh (University of Pittsburgh, 2010).

36. Erfani, R., Marefat, F. & Mohseni, P. Biosafety Considerations of a Capacitive Link for Wireless Power Transfer to Biomedical Implants. 2018 IEEE Biomed. Circuits Syst. Conf. BioCAS 2018 - Proc. 65, 923–927 (2018).

37. Erfani, R., Marefat, F., Sodagar, A. M. & Mohseni, P. Modeling and characterization of capacitive elements with tissue as dielectric material for wireless powering of neural implants. IEEE Trans. Neural Syst. Rehabil. Eng. 26, 1093–1099 (2018).

38. Sodagar, A. M. & Amiri, P. Capacitive coupling for power and data telemetry to implantable biomedical microsystems. 2009 4th Int. IEEE/EMBS Conf. Neural Eng. NER’09 411–414 (2009). doi:10.1109/NER.2009.5109320

39. Narayanamoorthi, R. Modeling of Capacitive Resonant Wireless Power and Data Transfer to Deep Biomedical Implants. IEEE Trans. Components, Packag. Manuf. Technol. 9, 1253–1263 (2019).

40. Mackenzie, J. W., Schuder, J. C. & Stephenson, H. E. Radio-frequency energy transport into the body. J. Surg. Res. 7, 133–144 (1967).

41. Erfani, R., Marefat, F., Sodagar, A. M. & Mohseni, P. Modeling and Experimental Validation of a Capacitive Link for Wireless Power Transfer to Biomedical Implants. IEEE Trans. Circuits Syst. II Express Briefs 65, 923–927 (2018).

42. International Commission on Non-Ionizing Radiation Protection. Guidelines for Limiting Exposure to Electromagnetic Fields (100 kHz to 300 GHz). Health physics 118, (2020).

43. Institute of Electrical and Electronics Engineers. IEEE Std C95.1™-2019: IEEE Standard for Safety Levels with Respect to Human Exposure to Electric, Magnetic, and Electromagnetic Fields, 0 Hz to 300 GHz. (Institute of Electrical and Electronics Engineers, 2019). doi:10.1109/IEEESTD.2019.8859679

44. Bureau Radiocommunication - International Telecommunication Union. Radio Regulations Articles Edition of 2020. (2020).

45. Weber, D. J. et al. BIONic WalkAide for correcting foot drop. IEEE Trans. Neural Syst. Rehabil. Eng. 13, 242–246 (2005).

46. Ausra, J. et al. Wireless battery free fully implantable multimodal recording and neuromodulation tools for songbirds. Nat. Commun. 12, 1–12 (2021).

47. Tudela-Pi, M., Minguillon, J., Becerra-Fajardo, L. & Ivorra, A. Volume Conduction for Powering Deeply Implanted Networks of Wireless Injectable Medical Devices: A Numerical Parametric Analysis. IEEE Access 9, 100594–100605 (2021).

48. Olivo, J., Carrara, S. & De Micheli, G. Biofuel cells and inductive powering as energy harvesting techniques for implantable sensors. Sci. Adv. Mater. 3, 420–425 (2011).

49. Heetderks, W. J. RF Powering of Millimeter- and Submillimeter-Sized Neural Prosthetic Implants. IEEE Trans. Biomed. Eng. 35, 323–327 (1988).

50. Becerra-Fajardo, L., Tudela-Pi, M. & Ivorra, A. Two-Port Networks to Model Galvanic Coupling for Intrabody Communications and Power Transfer to Implants. in IEEE Biomedical Circuits and Systems Conference (BioCAS) 20–23 (2018).

51. Carr, J. C., Richard Fright, W. & Beatson, R. K. Surface interpolation with radial basis functions for medical imaging. IEEE Trans. Med. Imaging 16, 96–107 (1997).

52. Field, D. A. Laplacian smoothing and Delaunay triangulations. Commun. Appl. Numer. Methods 4, 709–712 (1988).

53. Institute of Electrical and Electronics Engineers. IEC/IEEE Draft International Standard for Determining the Peak Spatial Average Specific Absorption Rate (SAR) in the Human Body from Wireless Communications Devices, 30 MHz - 6 GHz. Part 1: General Requirements for using the Finite Difference Time Domain. IEEE P62704-1D4, 2016 (2016).

