## Supplementary information for "Powering electronic implants by high frequency volume conduction: in human validation"

### Supplementary descriptions of the methods

#### Supplementary method 1: Marking procedure of electrodes' location for MRI acquisition and segmentation

Due to the incompatibility of the electrodes with MRI, a marking procedure was performed to accurately determine the positions of both the band electrodes (i.e. the external electrodes) and the needle electrodes for building the 3D computational models from the MRI images.

After the anatomical identification of the target muscle, the intended insertion point of the needle electrodes was marked by drawing a cross using a permanent ink marker. The edges of the bands were also marked. Eight MRI markers (PinPoint® for Small Field of View Imaging 187 from Beekley Corporation, Bristol, CT, US) were positioned on the ink markings: two MRI markers were used for the needle electrodes (i.e. one per needle) and three for each band. This arrangement allows to accurately obtain the position of the bands for building the 3D computational models. The marking procedure was performed for both limbs.

All the MRI markers were removed after the MRI acquisition, while the ink markings corresponding to the electrodes location were kept for the second phase of the experimentation. Supplementary figure 1 shows an example of the placement of some MRI markers on the arm of P1.

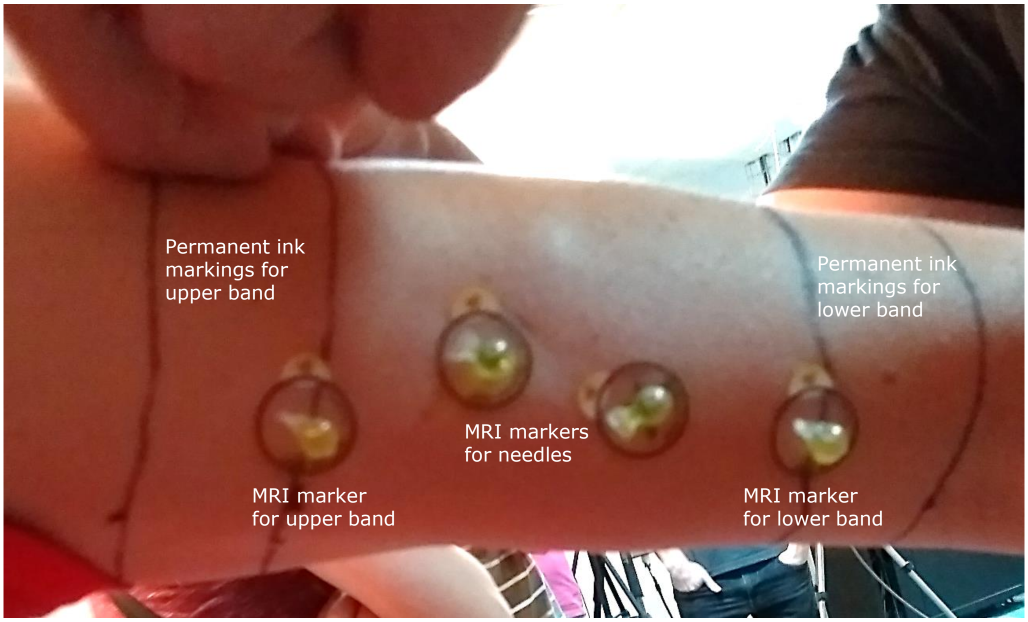


**Supplementary figure 1 | Picture of the placement of some MRI markers on the arm of P1.**

#### Supplementary method 2: HF voltage generator, applied waveform and external electrodes

**HF voltage generator.** The high frequency sinusoidal voltage bursts were generated using an arbitrary waveform generator (4065 from B&K Precision, Yorba Linda, CA, US). These bursts were amplified by using a custom-made class AB amplifier. This amplifier has a modular architecture and can provide, at 6.78 MHz, a maximum output voltage amplitude of approximately 200 V and a maximum output current amplitude of approximately 2.5 A. (Additional technical details regarding its architecture are intentionally omitted for preserving industrial property rights.) A picture of the custom-made amplifier is shown in Supplementary figure 2.

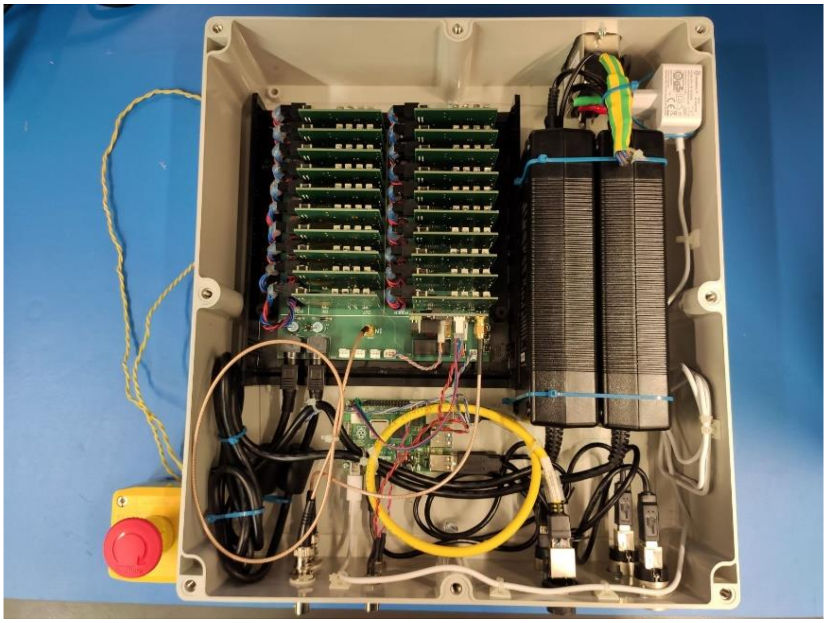

Supplementary figure 2 | Picture of the custom-made class AB amplifier. It has a modular architecture and can provide, at 6.78 MHz, a maximum output voltage amplitude of approximately 200 V and a maximum output current amplitude of approximately 2.5 A.

Applied waveform. To minimize the contribution of the harmonics generated by windowing the sinusoidal waveform, the applied bursts were smoothed with a tapered cosine window. This window is defined as

$$w(t) = \begin{cases} \frac{1}{2}\left[1-\cos\left(\frac{2\pi t}{rB}\right)\right], & 0 \le t < \frac{r}{2}B \\ 1, & \frac{r}{2}B \le t < B - \frac{r}{2}B \\ \frac{1}{2}\left[1-\cos\left(\frac{2\pi t}{rB} - \frac{2\pi}{r}\right)\right], & B - \frac{r}{2}B \le t < B, \end{cases} \quad \text{(S1)}$$

where $r$ is the cosine fraction and $B$ is the burst duration (i.e. window duration). This window is a rectangular window with the first and last $r/2$ percent of the time equal to parts of a cosine. The expression for a single smoothed burst with a sinusoid as the modulated waveform is

$$b(t) = v(t) \cdot w(t) = V_{peak} \cdot \sin(2\pi f t + \varphi) \cdot w(t), \quad \text{(S2)}$$

where $V_{peak}$, $f$ and $\varphi$ are the amplitude, frequency and initial phase of the sinusoidal modulated waveform respectively. Therefore, the RMS voltage of the waveform during the burst ($V_{rms\_in\_burst}$) is

$$V_{rms\_in\_burst} = \sqrt{\frac{1}{B}\int_0^B [v(t) \cdot w(t)]^2 dt}\,. \quad \text{(S3)}$$

If we assume $\varphi = 0$ and $rBf/2$ is an integer number higher than zero (which corresponds to our case), then

$$V_{rms\_in\_burst} = \frac{V_{peak}}{\sqrt{2}}\sqrt{1 - \frac{5}{8}r} = V_{peak} \cdot k, \quad \text{(S4)}$$

where $k$ is a constant that depends on the cosine fraction $r$. The RMS voltage of the waveform ($V_{rms}$) is

$$V_{rms} = V_{peak} \cdot k\sqrt{FB}, \quad \text{(S5)}$$

where $FB$ is the duty cycle, being $F$ and $B$ the repetition frequency and the duration of the bursts, respectively. Therefore, for power calculation, a scaling factor $k\sqrt{FB}$ has to be applied to transform amplitude values into RMS values for the applied waveform. A cosine fraction $r = 0.5$ was used, thus $k = \sqrt{11/32}$. A smoothed burst ($V_{peak}$ = 1 V, f = 6.78 MHz, B = 100 μs) is shown in Supplementary figure 3.

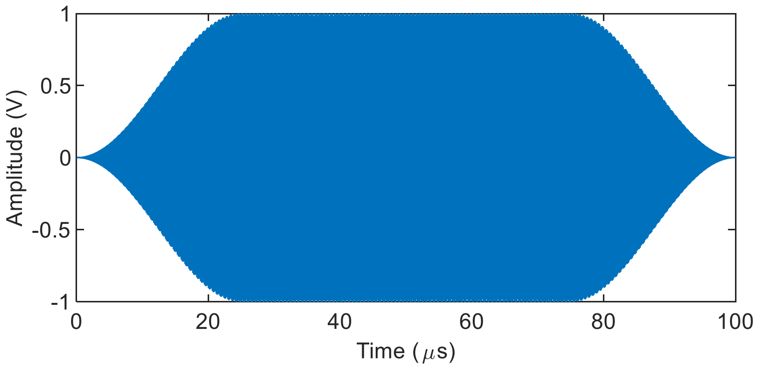


Supplementary figure 3 | Smoothed burst (Vpeak = 1 V, f = 6.78 MHz, B = 100 μs) with tapered cosine window (r = 0.5).

**External electrodes.** The external voltage was applied across a pair of external electrodes shaped as bands strapped around the limb and encompassing the region where the needle electrodes were inserted (see Supplementary figures 4 and 5). These electrodes were made with a conductive and biocompatible fabric based on silver-coated yarns (Shieldex® Technik-tex P130 + B from Statex Produktions und Vertriebs GmbH, Bremen, Germany).

### Supplementary method 3: Measurement apparatus

**Electrical measurements (ac).** The electrical ac measurements were acquired using a floating digital oscilloscope (TPS2014 from Tektronix, Inc., Beaverton, OR, US). For the applied external voltage and the potentiometer voltage, active differential probes (TA043 from Pico Technology Ltd, Saint Neots, UK) were used. A current probe (TCP2020A from Tektronix, Inc., Beaverton, OR, US) was used for the applied external current (i.e. current corresponding to the applied external voltage). This is shown in Supplementary figure 4.

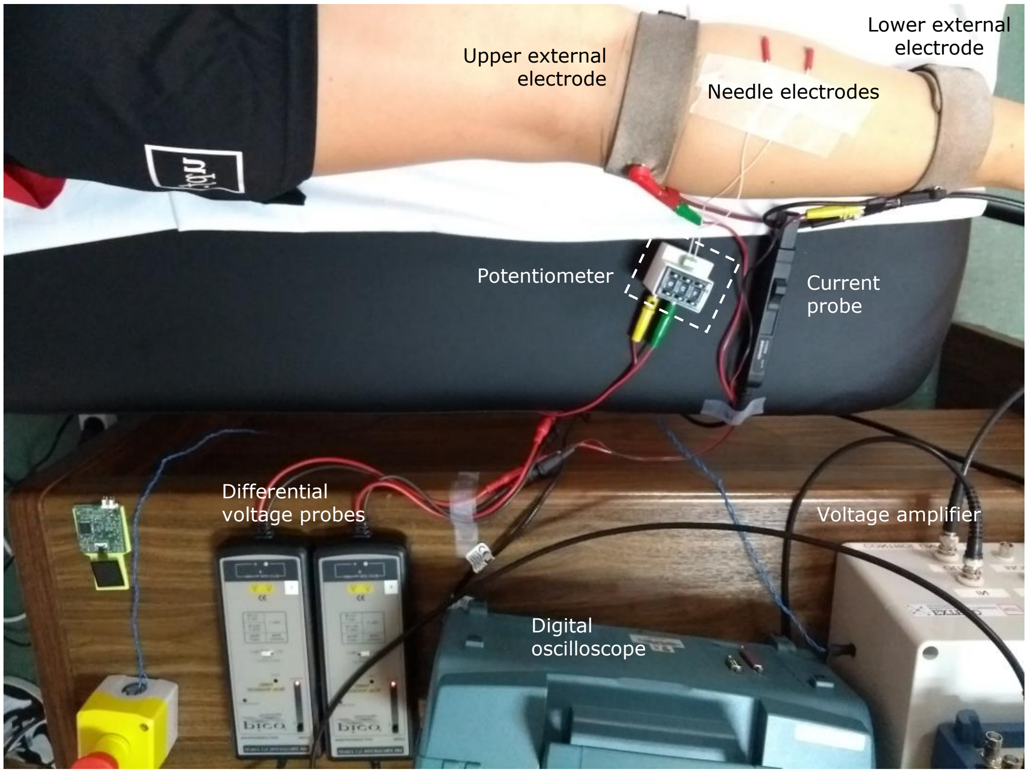


Supplementary figure 4 | Experimental ac measurement setup.

**Electrical measurements (dc).** The electrical dc measurements (i.e. voltage and current to calculate the power consumption of the demonstrative electronic device connected as load to the needle electrodes) were acquired using two multimeters (38XR-A from Amprobe, Everett, WA, US): one to measure voltage and another one to measure current. This is shown in Supplementary figure 5.

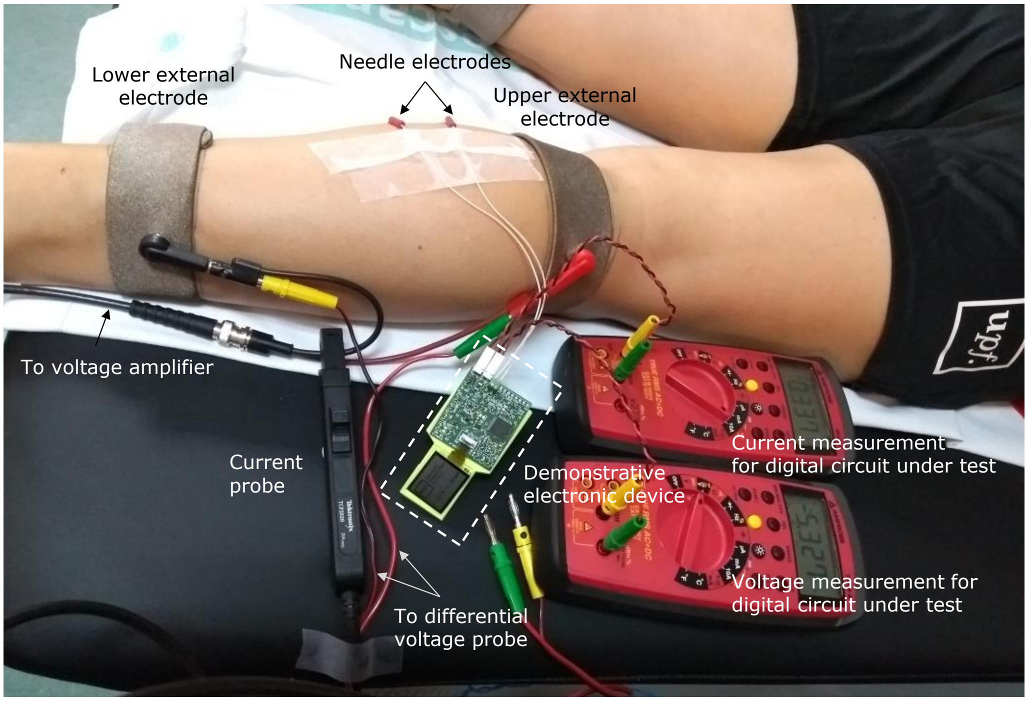


**Supplementary figure 5 | Experimental dc measurement setup.**

**Temperature measurements.** The temperature measurements were acquired and recorded using an infrared camera (E60 from FLIR Systems Ltd, Wilsonville, OR, US).

### Supplementary method 4: Demonstrative electronic device connected to needle electrodes

To illustrate the potential of volume conduction to power complex digital implants, a demonstrative circuit was designed and manufactured using commercial off-the-shelf components. This electronic device was connected to the needle electrodes. The main function of this circuit was to show its input voltage and current by means of a 1.3” LCD screen (LS013B7DH05 from Sharp Corporation Sakai, Osaka Prefecture, Japan).

The electronic device is composed of three subcircuits: 1) power stage, 2) sensing stage and 3) control unit and display. The power stage consists of a dc-blocking capacitor for each electrode, followed by a bridge rectifier (diode MCL103B from Vishay Intertechnology, Malvern, PA, US), a smoothing capacitor (10 μF) that rectifies the picked-up high frequency voltage, and a linear voltage regulator (TLV701 from Texas Instruments, Dallas, TX, US) that fixes a voltage of 3.3 V. Supplementary figure 6 shows the circuit architecture of this stage.

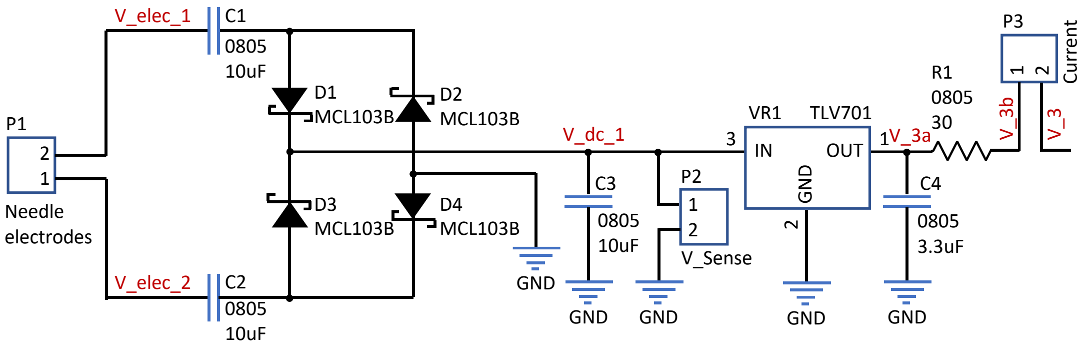


**Supplementary figure 6 | Power stage of the digital circuit.**

The sensing stage is used to measure the input dc voltage and current (see Supplementary figure 7). The voltage is measured across the smoothing capacitor that follows the diode bridge. The current is measured by acquiring the voltage drop across a shunt resistor (30 Ω) located after the smoothing capacitor of the regulator's output. This voltage is amplified using two amplification stages (implemented with operational amplifiers TLV521 from Texas Instruments, Dallas, TX, US) that set a gain of 51 V/V. The voltages are digitized using two 8-bit analog-to-digital converters (ADS7040 from Texas Instruments, Dallas, TX, US). The result of the conversions is obtained by the control unit through a serial peripheral interface (SPI).

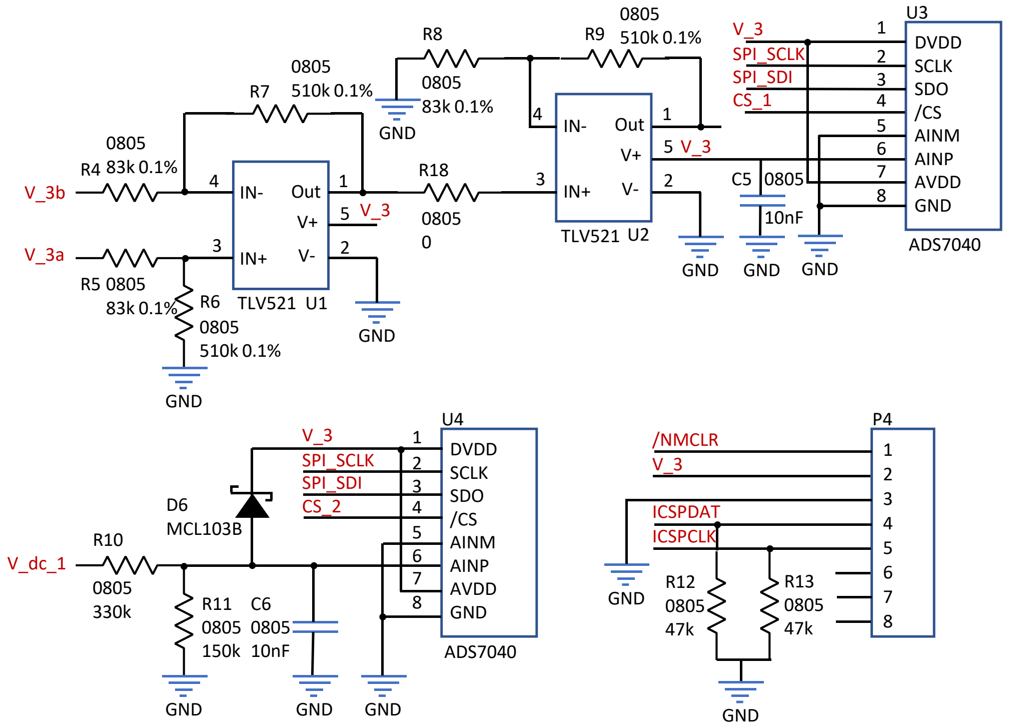


**Supplementary figure 7 | Sensing components of the designed circuit and programming connector (P4).**

The control unit is based on an 8-bit, low-power microcontroller (PIC18LF47K42 from Microchip Technology Inc., Chandler, AZ, US) with a clock frequency set to 1 MHz (see Supplementary figure 8). The microcontroller interrogates the ADCs every 10 ms and then updates the LCD screen (via SPI communication) with the acquired measurement.

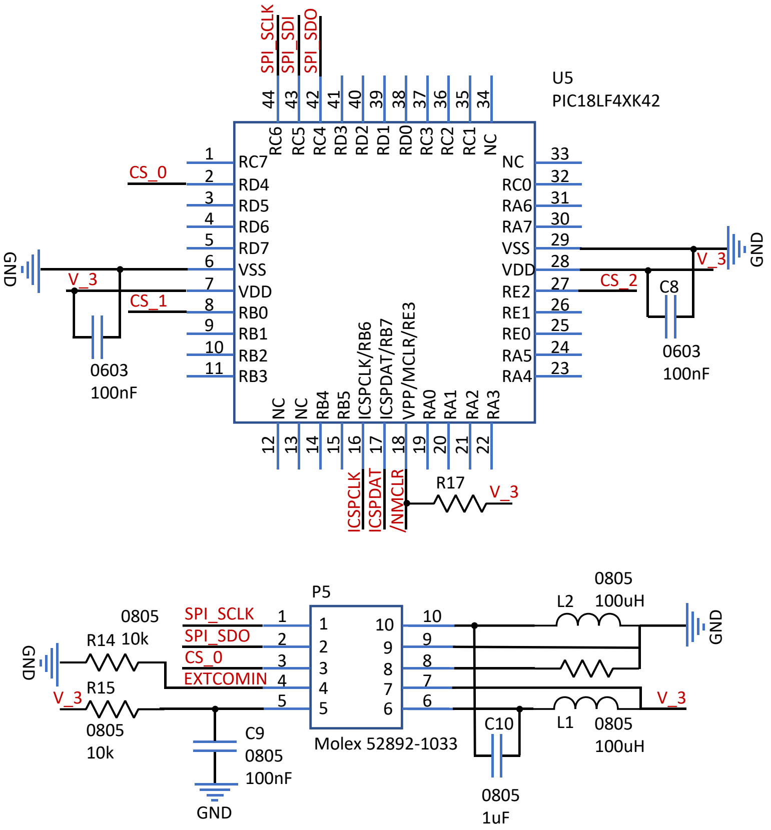


Supplementary figure 8 | Microcontroller and LCD interface connector (P5).

Both current (mA) and voltage (V) are shown on the screen with a resolution of 0.1 mA and 0.1 V, respectively (see Supplementary figure 9). Approximately every 20 s the microcontroller refreshes the LCD and changes the logo that the LCD shows on its top. Two different logos can be displayed. Since the main purpose of this device was to show the potential of volume conduction to power complex digital circuits, the circuit was designed to minimize its power consumption at the expense of the measurement accuracy. That is the reason why the voltage and the current to calculate the power consumption were externally measured through the two 2-pin connectors P2 and P3 using the multimeters mentioned in Supplementary method 3.

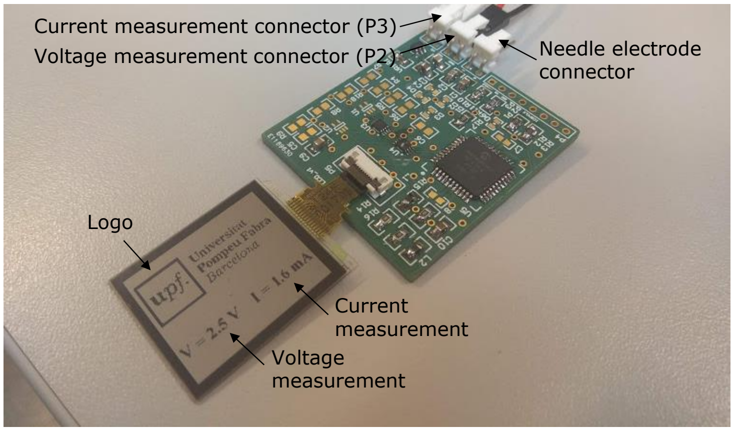


Supplementary figure 9 | Picture of the electronic device.

The electronic components were mounted on a 45 × 40 mm two-layers PCB (see Supplementary figure 10). The base material is FR-4 and its thickness is 1.55 mm.

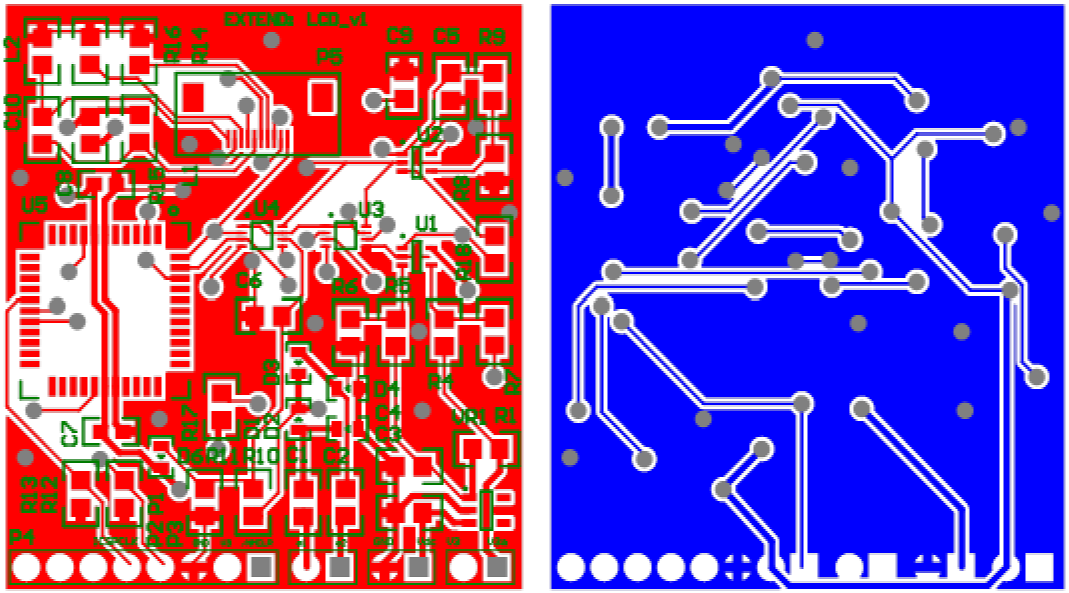

**Supplementary figure 10 | PCB layout.** Top layer (red), top overlay (green) and bottom layer (blue).

### Supplementary method 5: Numerical methods

The segmented meshes (from the MRI images) of the tissues were imported into COMSOL Multiphysics 5.3 (from COMSOL, Inc., Burlington, MA, US) to create the 3D computational model. To numerically determine the electric field and voltage distributions inside the tissues, the "electric current" physics that is included inside the ac/dc module of COMSOL Multiphysics 5.3 was used. The equation that this modelling software solves is the continuity equation of the electromagnetism

$$\nabla \cdot J = Q, \qquad \text{(S6)}$$

where $Q$ is the charge and the current density, $J$, is defined as

$$J = \sigma E + j\omega D + J_{\mathrm{e}}, \qquad \text{(S7)}$$

being $\sigma$ the conductivity, $\omega$ the angular frequency, $D$ the electric displacement field, $J_{\mathrm{e}}$ the external applied current density, and $E$ the electric field

$$E = -\nabla V, \qquad \text{(S8)}$$

where $V$ is the voltage.

The geometry of each one of the ten studied 3D computational models consisted in a four-tissue layered segmented limb obtained from the MRI images (see Subsection "MRI acquisition and segmentation" in "Methods"). The four layers, from the most peripheral to the most internal one were skin, fat, muscle, and cancellous bone. Two cylindrical electrodes with a diameter of 0.4 mm, and a total length of 20 mm emulated the needle electrodes that we used in the experimental part. Their length was divided into two longitudinal sections: 17 mm of insulating material and 3 mm of exposed surface at the tip of the needle. The position of these electrodes was determined using the coordinates of the MRI markers (see Supplementary method 1). They were perpendicularly aligned with the skin tissue and were inserted 17.5 mm inside the tissues. The position coordinates of the external electrodes were also identified in the MRI using three markers per electrode. These data were used to obtain a three-point plane. Following, a plane parallel to the previous one was created with a separation distance of 30 mm for the arms and 40 mm for the calf (i.e. the width of the band electrodes used during the experimentation). Then, the superficial tissue area encompassed between both planes was considered the area of the external electrodes. Finally, the whole limb was set inside a block that emulated the air. The size of this block was adjusted for each case to guarantee a minimum of 2 cm gap of air in any direction. Supplementary figure 11 shows the resultant geometry of the 3D computational model of the arm of participant P5.

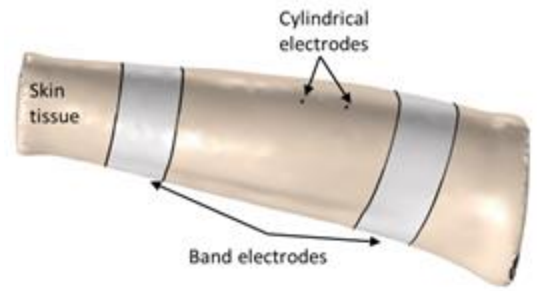


**Supplementary figure 11 | Geometry of the 3D computational model (COMSOL) of the arm of P5.** The air box that surrounded the arm has been intentionally hidden for visualization purposes.

Each one of the four tissues was assigned with its corresponding conductivity and permittivity at 6.78 MHz[1] and its mass density[1].These values are reported in Supplementary table 1. The electrical properties of the non-biological materials used for the simulation are reported in Supplementary table 2.

**Supplementary table 1 | Passive electrical properties and density of modeled tissues at a frequency of 6.78 MHz[1].**

| Dielectric properties @ 6.78 MHz | Conductivity (S/m) | Relative permittivity | Density (kg/m$^3$) |
|---|---|---|---|
| Skin | 0.147 | 478 | 1109 |
| Fat (subcutaneous) | 0.0496 | 35 | 911 |
| Muscle | 0.602 | 233 | 1090 |
| Bone (cancellous) | 0.116 | 90 | 1178 |

**Supplementary table 2 | Electrical properties of non-biological materials.**

| Dielectric properties | Conductivity (S/m) | Relative permittivity | Density (kg/m$^3$) |
|---|---|---|---|
| Conductive electrodes (Steel, 0.65% carbon) | $5.5\cdot10^{6}$ [2] | 1 | 7844 [2] |
| Insulating material of needle electrodes | $1\cdot10^{-17}$ [3] | 3.2 [3] | 1000 |
| Air (at 20º) | $1\cdot10^{-9}$ | 1 | 1.2 |

**Two-port model and Z-parameters extraction.** The electrical coupling between the external electrodes and the needle electrodes was modeled as a two-port network. This model included the external electrodes, the limb tissues, and the needle electrodes (emulated as cylinders). Since all its elements were passive, the system could be considered reciprocal[4]. Therefore, voltages and currents at the network could be expressed as

$$\begin{pmatrix} V_{ext} \\ V_{int} \end{pmatrix} = \begin{pmatrix} Z_{ext} & Z_{12} \\ Z_{12} & Z_{int} \end{pmatrix} \begin{pmatrix} I_{ext} \\ I_{int} \end{pmatrix}, \qquad \text{(S9)}$$

where $V_{ext}$ is the voltage across the external electrodes, $V_{int}$ is the voltage across the needle electrodes, $I_{ext}$ is the current through the two external electrodes and $I_{int}$ is the current through the needle electrodes. By definition,

$$Z_{ext} \stackrel{\text{def}}{=} \left.\frac{V_{ext}}{I_{ext}}\right|_{I_{int}=0}, \qquad \text{(S10)}$$

$$Z_{\text{int}} \stackrel{\text{def}}{=} \left.\frac{V_{\text{int}}}{I_{\text{int}}}\right|_{I_{\text{ext}}=0}, \tag{S11}$$

$$Z_{12} \stackrel{\text{def}}{=} \left.\frac{V_{\text{int}}}{I_{\text{ext}}}\right|_{I_{\text{int}}=0} = \left.\frac{V_{\text{ext}}}{I_{\text{int}}}\right|_{I_{\text{ext}}=0}. \tag{S12}$$

The impedances $Z_{\text{ext}}$ (equation (S10)) and $Z_{12}$ (equation (S12)) were determined by simulating the delivery of a reference current (1 A at 6.78 MHz) through the external electrodes, while keeping $I_{\text{int}}$ equal to 0, and measuring the voltage across the external electrodes (for $Z_{\text{ext}}$) and the voltage across the needle electrodes (for $Z_{12}$). The same procedure was done for determining $Z_{\text{int}}$ (equation (S11)) but, in this case, applying a reference current through the needle electrodes and measuring the voltage across the needle electrodes.

The parameters obtained for the five participants are summarized in Supplementary tables 3 and 4, for the arm and the lower leg respectively. Note that the impedance parameters $Z_{\text{int}}$ and $Z_{12}$ include three different angles (0º, -10º, +10º). As the needle electrodes were inserted manually, a misalignment of $\pm$ 10º between both electrodes could be obtained (see Figure 3a). An alignment of 0º corresponds to a parallel alignment between both cylindrical electrodes, -10º corresponds to the case when the distance between both tip electrodes was minimum and +10º corresponds to the case when the separation distance on the tips was maximum.

Supplementary table 3 | Arm impedance parameters at 6.78 MHz.

| Impedances | | P1 | P2 | P3 | P4 | P5 |
|---|---|---|---|---|---|---|
| $Z_{\text{ext}}$ (Ω) | | 94.1-29i | 95-28.2i | 104.4-31.6i | 58.8-15i | 56-19.2i |
| $Z_{\text{int}}$ (Ω) | 0 º | 423-63i | 460-69i | 432-65i | 420-62i | 416-63i |
| | -10 º | 418-62i | 454-68i | 424-63i | 411-61i | 411-62i |
| | +10 º | 430-64i | 484-73i | 444-67i | 431-64i | 420-64i |
| $Z_{12}$ (Ω) | 0 º | 18.7-3.9i | 20.1-4i | 18.9-4i | 13-2.3i | 13.2-3.2i |
| | -10 º | 15.2-3.1i | 16.9-3.4i | 14.7-3.1i | 8.1-1.5i | 10.6-2.5i |
| | +10 º | 22.2-4.6i | 23.4-4.7i | 23.1-4.9i | 17.8-3.2i | 15.8-3.8i |

Supplementary table 4 | Lower leg impedance parameters at 6.78 MHz.

| Impedance | | P1 | P2 | P3 | P4 | P5 |
|---|---|---|---|---|---|---|
| $Z_{\text{ext}}$(Ω) | | 79.2-19i | 72.3-20.1i | 86.9-23.7i | 60.2-15.1i | 61.3-15.5i |
| $Z_{\text{int}}$ (Ω) | 0 º | 429-63i | 423-63i | 491-73i | 418-62i | 419-62i |
| | -10 º | 425-63i | 420-62i | 484-72i | 419-62i | 415-61i |
| | +10 º | 438-65i | 428-63i | 547-83i | 410-60i | 424-63i |
| $Z_{12}$ (Ω) | 0 º | 7.6-1.3i | 7.7-1.3i | 6.7-1.2i | 7.6-1.3i | 7.5-1.3i |
| | -10 º | 6.4-1.1i | 6.3-1.1i | 5.5-1i | 6-1i | 6.1-1.1i |
| | +10 º | 8.9-1.5i | 9.1-1.6i | 8-1.4i | 9.3-1.6i | 8.9-1.6i |

The equivalent circuit for the Z-parameters of a reciprocal network is a T-circuit (see Supplementary figure 12). The power delivered to the load (PDL) can be expressed as

$$P_{\mathrm{Load}} = |I_{\mathrm{Load}}|^2 \Re(Z_{\mathrm{Load}}) \,, \tag{S13}$$

where $P_{\mathrm{Load}}$ is the power dissipated at the load, and $I_{\mathrm{Load}}$ is the current flowing through the load. $I_{\mathrm{Load}}$ can be calculated as

$$I_{\mathrm{Load}} = I_{\mathrm{ext}} \frac{Z_{12}}{Z_{\mathrm{int}} + Z_{\mathrm{Load}}}, \tag{S14}$$

being $I_{\mathrm{ext}}$ the current applied through the external electrodes. The power transmission efficiency (PTE) of the system is calculated as

$$PTE = \frac{P_{\mathrm{Load}}}{P_{\mathrm{Total}}} \cdot 100. \tag{S15}$$

where $P_{\mathrm{Total}}$ is the total power delivered through the external electrodes.

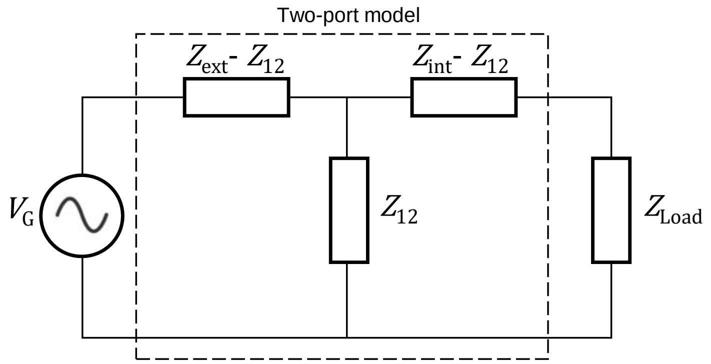


**Supplementary figure 12 | T-circuit used to determine the PDL and the PTE of the system.**

The load impedance ($Z_{Load}$) was the optimal load, or in other words, the resistance that experimentally maximized the received ac power (with a resolution of 10 Ω). The optimal loads for each participant and limb are reported in Supplementary table 5.

**Supplementary table 5 | Optimal $Z_{\mathrm{Load}}$ for each participant and limb.**

| $Z_{\mathrm{Load}}$ (Ω) | P1 | P2 | P3 | P4 | P5 |
|---|---|---|---|---|---|
| Arm | 210 | 250 | 260 | 250 | 250 |
| Lower leg | 260 | 250 | 280 | 250 | 250 |

**SAR calculation.** According to the IEEE safety standard[5], the specific absorption rate (SAR) can be calculated at any point by

$$SAR = \frac{\sigma |E_{rms}|^2}{\rho}, \tag{S16}$$

where $\sigma$ is the local tissue conductivity (S/m), $\rho$ is the local tissue density (kg/m$^3$) and $E_{rms}$ is the RMS value of the computed electric field (V/m). The SAR has to be is space-averaged over any cubical 10 g of tissue and time-averaged for 6 minutes using at each point equation (S16)[5].

The 3D computational model was used to simulate the electric field and the voltage distributions inside the tissues. For that, it was simulated the delivery of a sinusoidal voltage (frequency = 6.78 MHz) across the external electrodes, being its RMS value equal to the RMS value of the experimental applied waveform. The transformation from amplitude values into RMS values was done by applying the mentioned scaling factor $k\sqrt{FB}$, with $k = \sqrt{11/32}$ (see Supplementary method 2).

The resulting electric field and the voltage distributions were sampled using a 3D grid with a regular path of 1 mm. For each point, its coordinates, tissue conductivity, tissue density, and the obtained electric field and voltage were stored in a comma separated value (.csv) file. This file was imported to MATLAB R2019a (from Mathworks, Inc.,

Natick, MA, US). Afterwards, a moving voxel that consisted of a cube with a fixed edge of 21 mm (i.e. an averaged tissue mass value of approximately 10 g) and a path of 1 mm was defined as space-averaging tool. Subsequently, the voxel was swept through the whole imported geometry. For each voxel position, the space-averaged SAR was directly calculated as the mean of the punctual SAR (calculated from the electric field magnitude and equation (S16)) at any point within the voxel, and it was assigned to the central point of the voxel. The voxel positions that included any air point were not considered since the IEEE safety standard[5] indicate that the most peripheral parts of the extremities, which are in close contact with the air, can hardly have space-averaged thermal effects. Finally, the point of maximum space-averaged SAR was identified.

It must be noted that the SAR values reported in this work do not correspond to the actual SAR during the assays as in those assays the waveforms were applied for short periods (i.e. < 30 seconds) and the standard indicates that the SAR must be time-averaged for 6-minutes. The SAR values reported here would correspond to the case in which the waveforms were continuously applied for 6 minutes or more. For that reason, the SAR is referred as *projected SAR* in this work.

The projected SAR figures (i.e. Figure 2d and Supplementary figure 13) are geometrical longitudinal cuts that include their points of maximum projected SAR. In these figures, the applied sinusoidal voltage in the simulation was adjusted to obtain a maximum projected SAR of 10 W/kg. Once the point of maximum projected SAR was identified using the above procedure, the projected SAR was plotted for the remaining points in the same longitudinal plane. The points in this longitudinal plane that did not belong to any valid voxel position (i.e. the most peripheral points) were assigned with the maximum projected SAR value among the voxel positions where these points belong, as indicated by the IEEE[6].

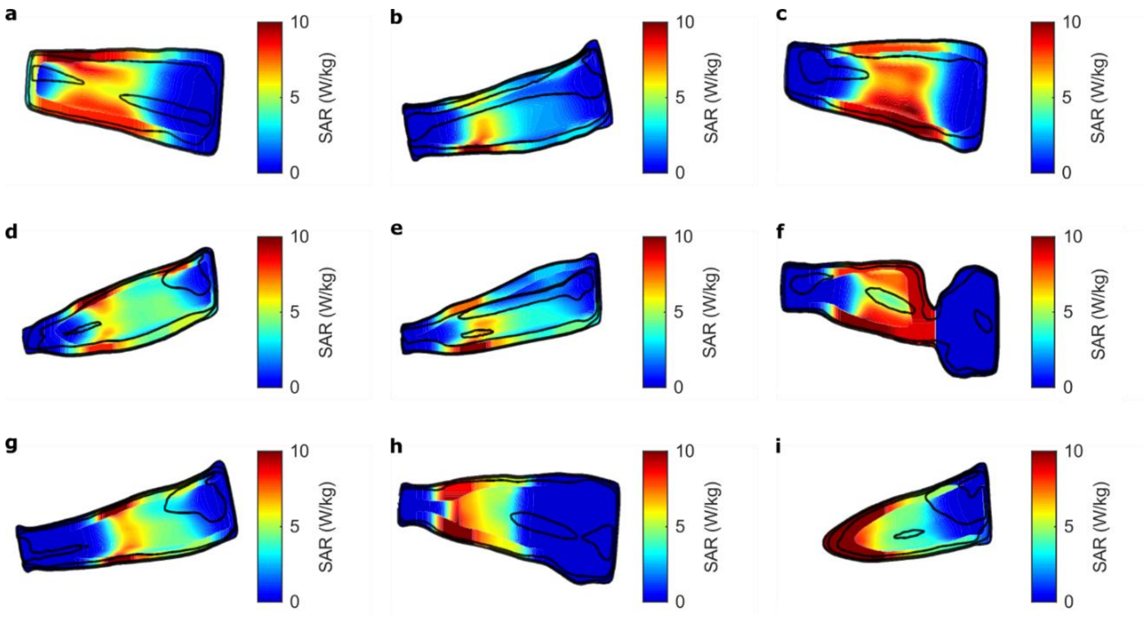


**Supplementary figure 13 | Computed projected SAR distribution for each case. a,** Arm of P1 (A1). **b,** Lower leg of P1 (LL1). **c,** Arm of P2 (A2). **d,** Lower leg of P2 (LL2). **e,** Lower leg P3 (LL3). **f,** Arm of P4 (A4). **g,** Lower leg of P4 (LL4). **h,** Arm of P5 (A5). **i,** Lower leg of P5 (LL5). The projected SAR distribution for the arm of P3 (A3) is reported in Figure 2d.

The computed electric field and voltage distributions, from which the projected SAR were computed, are presented for the longitudinal planes in Supplementary figures 14 and 15, respectively. The electric field and voltage were intentionally omitted on the air layers to improve the visualization on the tissues.

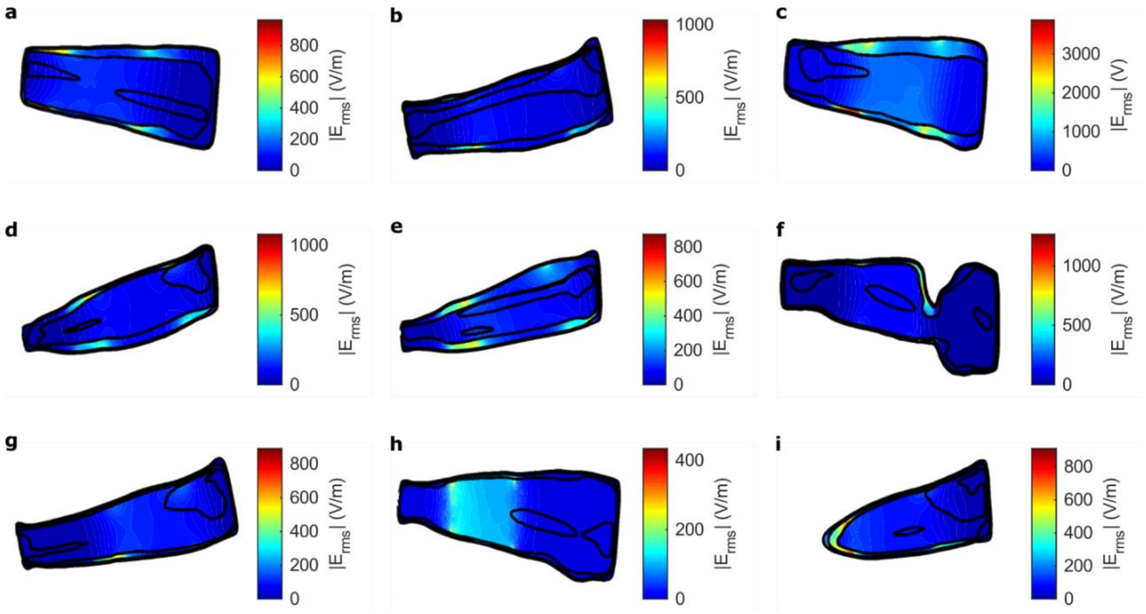


**Supplementary figure 14 | Computed electric field distribution for each case. a,** Arm of P1 (A1). **b,** Lower leg of P1 (LL1). **c,** Arm of P2 (A2). **d,** Lower leg of P2 (LL2). **e,** Lower leg of P3 (LL3). **f,** Arm of P4 (A4). **g,** Lower leg of P4 (LL4). **h,** Arm of P5 (A5). **i,** Lower leg of P5 (LL5). The electric field distribution for the arm of P3 (A3) is reported in Figure 2b.

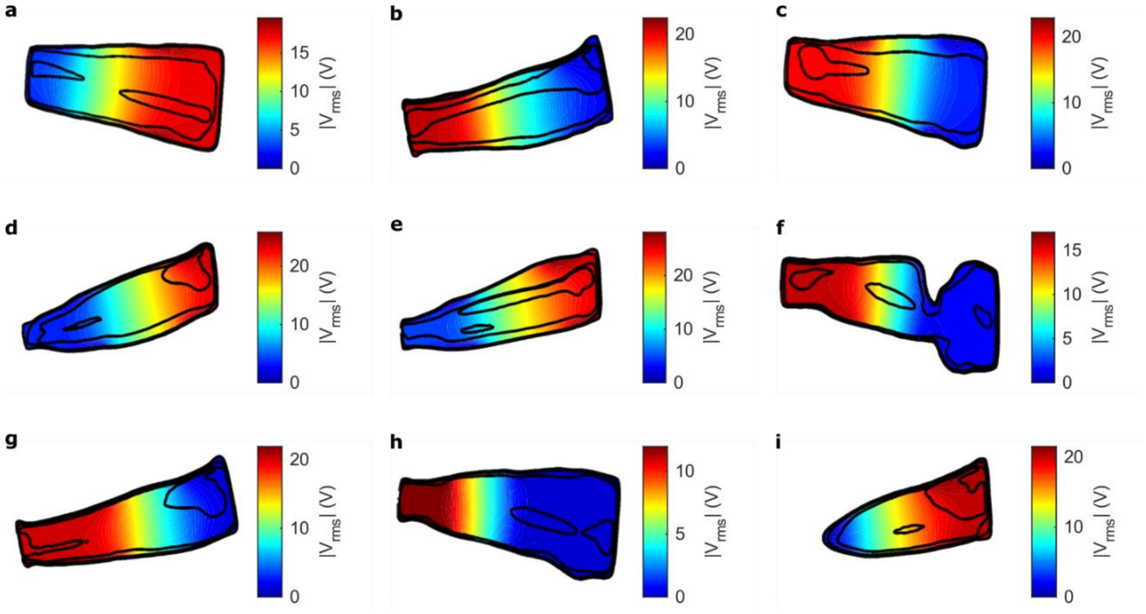


**Supplementary figure 15 | Computed voltage distribution for each case. a,** Arm of P1 (A1). **b,** Lower leg of P1 (LL1). **c,** Arm of P2 (A2). **d,** Lower leg of P2 (LL2). **e,** Lower leg of P3 (LL3). **f,** Arm of P4 (A4). **g,** Lower leg of P4 (LL4). **h,** Arm of P5 (A5). **i,** Lower leg of P5 (LL5). The voltage distribution for the arm of P3 (A3) is reported in Figure 2c.

**Maximum *in situ* electric field to avoid electrostimulation.** In terms of the electrostimulation effects, the IEEE safety standard[5] defines the dosimetric reference level (DRL) as the *in situ* electric field, and it is determined for frequencies between 0 Hz and 5 MHz. According to this, for continuous 6.78 MHz sinusoidal waveform, which is the carrier frequency of the sinusoidal voltage bursts used in this study, the DRL for electrostimulation mechanisms does not apply. However, the standard also provides limits for nonsinusoidal fields, as it is the case here. The standard indicates that the exposure waveform consisting of multiple frequencies must satisfy that

$$\sum_{0\,MHz}^{5\,MHz} \frac{A_i}{RL_i} \le 1, \tag{S17}$$

where $A_i$ is the magnitude of the $ith$ Fourier component of the sinusoidal voltage bursts, and $RL_i$ represents the *in situ* electric field restriction defined by the maximum allowed *in situ* electric field $E_i$ for the $ith$ Fourier component

$$E_i = \begin{cases} E_0 & for \quad f < f_e \\ E_0 \dfrac{f}{f_e} & for \quad f \ge f_e \end{cases}, \tag{S18}$$

where $E_0$ is the rheobase *in situ* field, $f_e$ is the transition frequency, and $f$ is the frequency of the $ith$ Fourier component. For the scenario considered in this study, in which the sinusoidal voltage bursts are applied to the limbs, $E_0$ is defined as 2.10 $V_{rms}$/m, and $f_e$ is defined as 3350 Hz.

From equations (S17) and (S18), two ranges are identified: a lower bound ($f \le$ 3350 Hz) and an upper bound (5 MHz $\ge f >$ 3350 Hz). Therefore,

$$\left( \sum_{0}^{3350\,Hz} \frac{A_i}{E_0} + \sum_{3350\,Hz}^{5\,MHz} \frac{A_i}{E_0 \frac{f_i}{f_e}} \right) \le 1. \tag{S19}$$

The maximum peak electric field that could be applied by the voltage generator was calculated according to equation (S19) using MATLAB R2019a. The sinusoidal voltage burst waveform (carrier frequency = 6.78 MHz, burst duration = 100 µs and repetition frequency = 1 kHz) was smoothed with a tapered cosine window created using the *tukeywin* function from MATLAB, with r = 0.5. A 0.1 s duration waveform was then generated using a sampling frequency of 100 Msps, and its discrete Fourier transform was calculated. After identifying the two ranges defined by equation (S19), their summation was computed having in mind the magnitude of the $ith$ Fourier component, the rheobase *in situ* field, and the transition frequency. The resulting maximum peak electric field (i.e. the maximum *in situ* electric field to avoid electrostimulation) was approximately 227 MV/m.

### Additional supplementary figures and tables

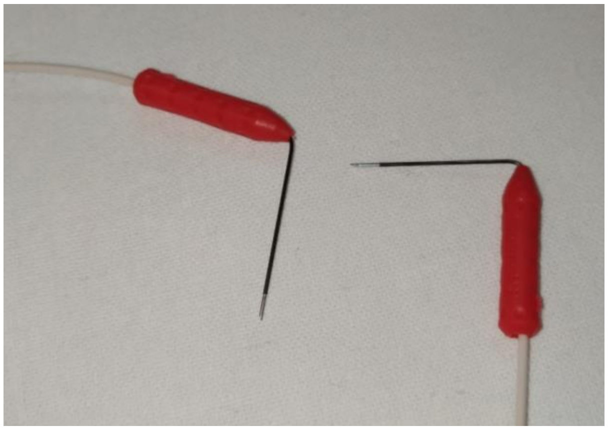

**Supplementary figure 16 | Picture of the needle electrodes after extraction.** No damage was observed in the electrodes.

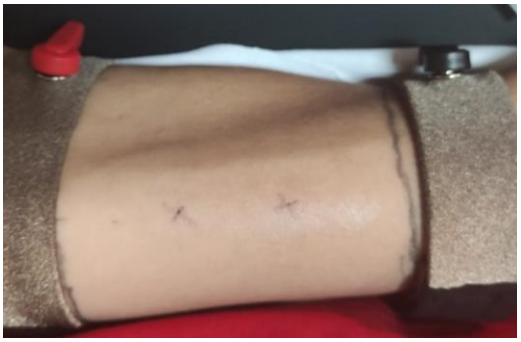

**Supplementary figure 17 | Picture of the arm of P4 after the extraction of the needles.** No damage was observed.

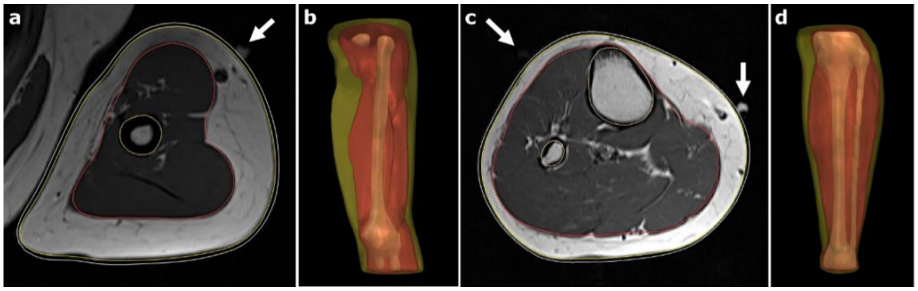

**Supplementary figure 18 | Examples of smoothed surface meshes and markers position from MRI images. a,** Axial slice of arm. **b,** 3D representation of segmented arm. **c,** Axial slice of lower leg. **d,** 3D representation of segmented lower leg. Bone mesh in brown, muscle in red, fat in yellow, and skin in white (only in a and c). White arrows indicate the MRI markers for electrode placement.

**Supplementary table 6 | Measured skin thickness from MRI images for each participant and limb.**

| Skin thickness (mm) | P1 | P2 | P3 | P4 | P5 |
|---|---|---|---|---|---|
| Arm | 1.75 | 1.50 | 1.50 | 2.00 | 1.75 |
| Lower leg | 1.50 | 2.00 | 1.75 | 2.00 | 2.25 |

### References of Supplementary information


1. Hasgall, P. *et al.* IT'IS Database for thermal and electromagnetic parameters of biological tissues, Version 4.0. *IT'IS* (2018).
2. Physical Data on the Elements and Alloys. in *Fundamentals of Modeling for Metals Processing* (eds. Furrer, D. U. & Semiatin, S. L.) **22A**, (ASM International, 2009).
3. M, A. U., Anthony, E. & Godspower, E. Electrical Properties of Enamel Wire Insulation. *Int. J. Trend Sci. Res. Dev.* **3**, 803–806 (2019).
4. Becerra-Fajardo, L., Tudela-Pi, M. & Ivorra, A. Two-Port Networks to Model Galvanic Coupling for Intrabody Communications and Power Transfer to Implants. in *IEEE Biomedical Circuits and Systems Conference (BioCAS)* 20–23 (2018).
5. Institute of Electrical and Electronics Engineers. *IEEE Std C95.1™-2019: IEEE Standard for Safety Levels with Respect to Human Exposure to Electric, Magnetic, and Electromagnetic Fields, 0 Hz to 300 GHz*. (Institute of Electrical and Electronics Engineers, 2019). doi:10.1109/IEEESTD.2019.8859679
6. Institute of Electrical and Electronics Engineers. *IEC/IEEE Draft International Standard for Determining the Peak Spatial Average Specific Absorption Rate (SAR) in the Human Body from Wireless Communications Devices, 30 MHz - 6 GHz. Part 1: General Requirements for using the Finite Difference Time Domain. IEEE P62704-1D4, 2016* (2016).
