## Supplementary material (MATLAB scripts) for "Powering electronic implants by high frequency volume conduction: in human validation": MPE_calculation.m

```
% MATLAB scripts for computing the maximum in situ electric field to avoid 
% electrostimulation of the IEEE safety standard

Epeak = 1000; %V/m
f= 6.78e6; %Frequency of carrier
F = 1000; %Hz
B = 100e-6; %s
FB=F*B;

Limb_fe = 3350; %Hz, According to IEEE standard
Eo = 2.1; %Persons permitted in restricted environments E0 (Vrms/m) 

Fs=100e6;
time_window=0.1;
t=(0:1/Fs:time_window-(1/Fs));
freqs=[0:Fs/size(t,2):(Fs)-(Fs/size(t,2))];

% Construct vector for sinusoidal bursts: normal or smoothed with tapered cosine (r=0.5)
    % y=Epeak*sin(2*pi*f*t).*(0.5*(square((2*pi*F*t),100*FB)+1)); %Normal waveform
    y = Epeak*sin(2*pi*f*t).*Tapered_cosine(B, Fs, F, t); %Smoothed waveform with tapered cosine r = 0.5

figure, plot(t,y), title('Original signal'), xlabel('Time');
z=fft(y)/(0.5*size(t,2));
figure, plot(freqs,abs(z)), title('FFT'), xlabel('Frequency');

indice_3350Hz=find(freqs==Limb_fe); %Find location in vector where 3350 Hz is located
indice_5MHz=find(freqs==5E6); %Find location in vector where 5 MHz is located

%Calculate value for lower bound: up to 3350 Hz
A1_lowerbound = abs(z(1:indice_3350Hz)); 
RL_lowerbound = Eo*sqrt(2);
sum1 = sum(A1_lowerbound/RL_lowerbound);
%Calculate values for upper bound: uo to 5 MHz
A1_upperbound = abs(z(indice_3350Hz+1:indice_5MHz));
RL_upperbound = Eo*sqrt(2)*freqs(indice_3350Hz+1:indice_5MHz)/Limb_fe;
sum2 = sum(A1_upperbound./RL_upperbound);
%Do the complete summation = lower bound + upper bound
complete_summation=sum1+sum2;

max_Epeak=Epeak/complete_summation

%% Verification
%Verify the maximum Epeak (max_Epeak) calculated above
clearvars -except max_Epeak f t FB Limb_fe Eo F freqs B Fs

% Construct vector for sinusoidal bursts: normal or smoothed with tapered cosine (r=0.5)
    % y=max_Epeak*sin(2*pi*f*t).*(0.5*(square((2*pi*F*t),100*FB)+1)); %Normal waveform 
    y = max_Epeak*sin(2*pi*f*t).*Tapered_cosine(B, Fs, F, t); %Tapered cosine r = 0.5

z=fft(y)/(0.5*size(t,2));
indice_3350Hz=find(freqs==Limb_fe);
indice_5MHz=find(freqs==5E6);
A1_lowerbound = abs(z(1:indice_3350Hz)); %Up to 3350 Hz
RL_lowerbound = Eo*sqrt(2);
sum1 = sum(A1_lowerbound/RL_lowerbound);
A1_upperbound = abs(z(indice_3350Hz+1:indice_5MHz)); %From 3350 Hz to 5 MHz
RL_upperbound = Eo*sqrt(2)*freqs(indice_3350Hz+1:indice_5MHz)/Limb_fe;
sum2 = sum(A1_upperbound./RL_upperbound);
complete_summation=sum1+sum2;
```
