## Supplementary material (MATLAB scripts) for "Powering electronic implants by high frequency volume conduction: in human validation": SAR_calculation.m

```
% MATLAB script for computing the SAR 

%csv file witht the stored results
H = csvread('S2_upper_limb.csv');

%tissue densities
rho_bone = 1178;
rho_muscle = 1090;
rho_fat = 911;
rho_skin = 1109;

x_v = H(:,1); %x-coordinate
y_v = H(:,2); %y-coordinate
z_v = H(:,3); %z-coordinate
E_v = H(:,4); %rms electric field
s_v = H(:,5); %tissue conductivity
v_v = H(:,6); %voltage

cx = 170; %londitudinal size
cy = 150; %transverse size
cz = 350; %depth size

%geometry reconstruction
x = reshape(x_v,cx,cy,cz);
y = reshape(y_v,cx,cy,cz);
z = reshape(z_v,cx,cy,cz);
E = reshape(E_v,cx,cy,cz);
s = reshape(s_v,cx,cy,cz);
v = reshape(v_v,cx,cy,cz);

clear x y z x_v y_v z_v E_v s_v s_v H


%conductivity
sigma_bone = 0.116;
sigma_muscle = 0.602;
sigma_fat = 0.0496;
sigma_skin = 0.147;


SAR = -50*ones(cy,cz); %%SAR matrix initalization
inh = 0;
max_sar = 0;
%swept through geometry
for x =80:1:(82)
    E_21 = E((x-10):1:(x+10),:,:);
    S_21 = s((x-10):1:(x+10),:,:);
    for  z = 11:1:(cz-10)
        for y = 11:1:(cy-10)
                sum_SAR = 0;
                    for y1 = -10:1:10
                        for x1 = -10:1:10
                            for z1 = -10:1:10
                                %if any point is air the sar is not
                                %averaged for this voxel
                                if S_21(11+y1,y+x1,z+z1) == 1E-9
                                    inh = 1;
                                else
                                % density in function of the tissue
                                    if S_21(11+y1,y+x1,z+z1) == sigma_muscle
                                        rho = rho_muscle;
                                    elseif S_21(11+y1,y+x1,z+z1) == sigma_bone
                                        rho = rho_bone;
                                    elseif S_21(11+y1,y+x1,z+z1) == sigma_fat
                                        rho = rho_fat;                                
                                    elseif S_21(11+y1,y+x1,z+z1) == sigma_skin
                                        rho = rho_skin;
                                    else
                                        rho = 1000;
                                    end
                                    %puncutal sar accumulation
                                    sum_SAR = sum_SAR + E_21(11+y1,y+x1,z+z1)*E_21(11+y1,y+x1,z+z1)*S_21(11+y1,y+x1,z+z1)/rho;
                                end
                            end
                        end
                    end
                
                if inh == 1
                    SAR(y,z)= -50;
                    inh = 0;
                else
                    %sar averaging for the validated voxels
                    SAR(y,z) = sum_SAR/(21*21*21);
                end
        end
    end
    %sar maximum
    if (max_sar < max(max(SAR)))
       max_sar = max(max(SAR));
       centre = x;
       copia_SAR = SAR;
    end
end

%21 mm slide
E_21 = E((centre-10):1:(centre+10),:,:);
v_21 = reshape(abs(v(centre,:,:)),cy,cz);
S_21 = s((centre-10):1:(centre+10),:,:);
SAR = copia_SAR;

%swept through the tissue points that couldnt be a center of a valid voxel
%in the previous step
for  z = 21:1:(cz-20)
    for x = 21:1:(cy-20)
        % air point
        if (copia_SAR(x,z) == -50 && S_21(11,x,z) == 1E-9)
            SAR(x,z) = -1; %the sar is assigned to a negative value for this case
        elseif (copia_SAR(x,z) == -50 && S_21(11,x,z) ~= 1E-9)
            %search to determine the maximum voxel where this point was
            %used
            max_SAR = -50;
            for x1 = -20:1:20
                for z1 = -20:1:20
                    if copia_SAR(x+x1,z+z1) > max_SAR &&  S_21(11,x+x1,z+z1) ~= 1E-9
                        max_SAR = copia_SAR(x+x1,z+z1);
                        if max_SAR == -50
                            max_SAR = -1;
                        end
                    end
                end   
            end
            SAR(x,z) = max_SAR;
        else
            SAR(x,z) = copia_SAR(x,z);
        end
    end
end


%simulation that includes the tissue density with a fixed step of 0.5 mm
H = csvread('S2_upper_limb_resol.csv');
s_v = H(:,5);
s2 = reshape(s_v,cx,cy*2,cz*2);
profile = reshape(s2(centre,:,:),cy*2,cz*2);      

%tissue edge detector
BW1 = (edge(profile,'Canny'));
imshow(((BW1)))
[row,col] = find(BW1);    
    
jet1 = [1 1 1; jet];

%SAR figure
xlabel('z coordinate (mm)');
ylabel('x coordinate(mm)');
[C,h] = contourf((SAR),200)
colormap(jet1)
set(h,'LineColor','none'); hold on;
plot1 = scatter(col/2,row/2,0.2,'o','MarkerFaceColor',[0 0 0],'MarkerEdgeColor',[0 0 0],'MarkerEdgeAlpha',0.2,'MarkerFaceAlpha',0.2); 
plot1.MarkerFaceAlpha = 0.2; hold on;
c = colorbar;
c.Label.String = 'SAR (W/kg)';
c.Limits = [0 10];
c.FontSize = 10;
c.FontName = 'Arial';
set(gca,'visible','off')
max(max(SAR))

%V figure
figure
[C,h] = contourf(real(v_21),200)
colormap(jet1)
set(h,'LineColor','none'); hold on;
plot1 = scatter(col/2,row/2,0.5,'o','MarkerFaceColor',[0 0 0],'MarkerEdgeColor',[0 0 0],'MarkerEdgeAlpha',0.2,'MarkerFaceAlpha',0.2); 
plot1.MarkerFaceAlpha = 0.2; hold on;
c = colorbar;
c.Label.String = '|V_{rms}| (V)';
c.Limits = [0 max(max(v_21))];
c.FontSize = 10;
c.FontName = 'Arial';
set(gca,'visible','off')

%E figure
figure
[C,h] = contourf((e21),200)
colormap(jet1)
set(h,'LineColor','none'); hold on;
plot1 = scatter(col/2,row/2,0.5,'o','MarkerFaceColor',[0 0 0],'MarkerEdgeColor',[0 0 0],'MarkerEdgeAlpha',0.2,'MarkerFaceAlpha',0.2); 
plot1.MarkerFaceAlpha = 0.2; hold on;
c = colorbar;
c.Label.String = '|E_{rms}| (V)';
c.Limits = [0 max(max(e21))];
c.FontSize = 10;
c.FontName = 'Arial';
set(gca,'visible','off')


 
```
