## Supplementary material (MATLAB scripts) for "Powering electronic implants by high frequency volume conduction: in human validation": Tapered_cosine.m

```
function [Tapered_cosine_window] = Tapered_cosine(B, Fs, F, t)
    Tapered_cosine = tukeywin(B*Fs);  % r = 0.5
    Burst = ([Tapered_cosine; zeros(((1/F)-B)*Fs,1)]);
    Repetitions = round(length(t)/length(Burst));
    Tapered_cosine_window = repmat(Burst,[Repetitions 1]).'; 
end
```
